# Proofreading and single-molecule sensitivity in T-cell receptor signaling by condensate nucleation

**DOI:** 10.1101/2024.03.13.584745

**Authors:** William L White, Hailemikael K Yirdaw, Ariel J Ben-Sasson, Jay T Groves, David Baker, Hao Yuan Kueh

**Affiliations:** Department of Bioengineering, University of Washington, Seattle, WA, USA; Institute for Protein Design, University of Washington, Seattle, WA, USA; Department of Chemistry, University of California Berkeley, Berkeley, CA, USA; Institute for Digital Molecular Analytics and Science, Nanyang Technological University, Singapore, Singapore; Department of Biochemistry, University of Washington, Seattle, WA, USA; Howard Hughes Medical Institute, Seattle, WA, USA; Institute for Stem Cell and Regenerative Medicine, University of Washington, Seattle, WA, USA

**Keywords:** T-cell receptor signaling, phase separation, nucleation-condensation, kinetic proofreading, stochastic computational modeling

## Abstract

T-cells display the remarkable ability to detect single foreign peptides displayed on target cells, while ignoring highly abundant self peptides. This selectivity has been explained by kinetic proofreading in the T-cell receptor (TCR) signaling pathway, which prevents responses to short-lived binding events regardless of their abundance. However, the biochemical mechanisms that drive kinetic proofreading have remained unclear. Here, using computational modeling, we show that these key signaling properties of the TCR pathway can emerge from the dynamics of LAT phosphorylation, diffusion, and condensation following TCR-pMHC binding. In this model, time delays in LAT condensate nucleation underlie kinetic proofreading, enabling selective signaling responses to high-affinity pMHC ligands. The cooperativity in the nucleation and growth of LAT condensates also provides a mechanism to amplify weak signals from single foreign peptides and for condensates to grow with increasing antigen numbers. In contrast to other models, condensate-nucleation proofreading predicts a dependence of signal strength on pMHC spacing at fixed number, a prediction we validated experimentally using a protein scaffold to present pMHCs at defined intervals. Our results suggest that nucleation-condensation proofreading underlies the remarkable antigen detection capabilities of the TCR signaling pathway.

**Significance:** To fight infections and cancer, T-cells must selectively recognize low levels of foreign peptides from pathogens or cancer cells, but the mechanisms that enable these properties have remained unclear.Using mathematical modeling and experiments, we find that T-cells can selectively detect single foreign peptides through a clustering process where a key protein downstream of the T-cell receptor forms condensates containing hundreds of signaling proteins. This condensate nucleation process can explain experimentally observed features of T-cell signaling, including our finding that the size of signaling clusters depends on the distance between foreign peptides. Our work reveals a key role for the condensation of signaling molecules in setting the spatial and temporal thresholds that control the sensitivity and selectivity of T-cells.

## Introduction

The TCR signaling pathway simultaneously achieves several functional properties that enable T-cells to sense and appropriately respond to challenges of varying types and severities^1^. Firstly, it senses foreign peptide antigens displayed on major histocompatibility complex ligands (pMHCs) in a highly sensitive manner, in some cases detecting single copies of foreign pMHCs^2^. Secondly, it is highly selective so as to avoid responding to more abundant self antigens, which have a weaker binding affinity for the TCR^3^. And finally, although the TCR signaling pathway activates in a digital, all-or-none manner upon pMHC recognition^4–6^, it can also tune its response to different levels of antigen input in an analog manner, enabling T-cells to tailor their responses to the magnitude of the threat^7,8^. Despite significant advances in our knowledge of the biochemical steps involved in TCR signaling^9,10^, it remains incompletely understood how the TCR signaling pathway is able to simultaneously achieve sensitivity, selectivity, and dynamic range in antigen sensing.

The first step in T-cell activation after a pMHC binds to the TCR is phosphorylation of the ITAM-containing cytoplasmic tails of the TCR-associated CD3 proteins by Lck^9^. The kinase ZAP70 can then bind to the phospho-ITAMs and become activated through phosphorylation by Lck and by trans-autophosphorylation^11^, and can in turn phosphorylate a variety of downstream substrates including the transmembrane protein LAT^9^, leading to signaling pathway activation. To explain how the T-cell signaling pathway is able to selectively respond to foreign peptides based on their stronger binding affinity, the concept of kinetic proofreading has been proposed^12^ and extensively studied using mathematical modeling^4,7,13–16^. In its most basic form, kinetic proofreading consists of a series of biochemical steps that can occur only when a TCR is bound to a pMHC, and that are quickly reversed when the pMHC unbinds^12^. This allows the T-cell to respond strongly to foreign antigens that bind long enough to complete all necessary biochemical steps, while ignoring self antigens that bind only briefly.

A number of extensions of the basic kinetic proofreading model have been evaluated, and shown to improve different facets of the signaling behavior. The addition of a negative feedback loop can dramatically enhance signaling selectivity^13^, whereas positive feedback loops can enhance sensitivity to low pMHC copy numbers^4,17^, generating an all-or-none ‘digital’ response. On the other hand, incoherent feedforward loops can expand the dynamic range of the T-cell response^7^. Together these models demonstrate each of the three critical signaling properties of T-cells, but individually each model is only able to achieve one or two. Additionally, most current models do not account for the inherent stochasticity in signaling which can result in responses that differ significantly from deterministic predictions^16,18^, and can dramatically reduce selectivity^19^.

The condensation and clustering of signaling molecules downstream of the TCR could constitute important events for recapitulating the cardinal properties of T cell signaling. Pioneering studies from Su and co-workers established that LAT can undergo condensation when phosphorylated. Phospho-tyrosine (pY) sites on phospho-LAT (pLAT) act as binding sites for SH2 domains in Grb2 and PLCγ, among other proteins^20,21^. Both Grb2 and PLCγ also contain SH3 domains that can bind to the proline-rich repeat regions of SOS^20–22^. This network of polyvalent interactions causes these molecules to form condensates tethered to the membrane by LAT^20–22^, and recruits a variety of other molecules which act together with PLCγ and SOS to transmit the activation signal into the T-cell^9,23^. These clustering events have been overlooked by existing kinetic proofreading models, which generally assume that the reaction system is well-mixed, even though it has been determined experimentally that this is not the case^20–22,24^. More recent work has explored the condensation of LAT with Grb2, SOS, and PLCγ using coarse grained molecular dynamics simulations^21,25^. However, these studies focus only on the formation of the condensate and not its context within the larger signaling pathway.

To address these gaps, we developed a computational model to study the dynamics of LAT condensation driven by TCR signaling. In this model, TCR antigen engagement leads to localized LAT phosphorylation and condensation at the cell membrane through a stochastic reaction-diffusion process. We find that this model shows all critical signaling properties of the TCR pathway, including its sensitivity to low ligand numbers, selectivity to high affinity ligands, and dynamic range in response to variations in pMHC ligand numbers. We also test key experimental predictions of this model, including activation dynamics, which provide a direct benchmark of the dynamics of pLAT cluster formation in response to single pMHC-TCR binding events^24^. Finally, we experimentally validate the spatial component of our model using a designed protein system^26^ that allows systematic variation of antigen spacing independent of antigen number. Together, these findings establish a quantitative framework for understanding the exceptional functional capabilities of the T cell in threat sensing and discrimination.

## Results

### The LAT condensation model

To assess the contribution of LAT condensation to TCR signaling we built a dynamical model to capture the spatial distributions of LAT at the membrane due to clustering. To achieve this property, we built our model on a 2D grid representing the plasma membrane of a T-cell. Within this 2D grid, we model the location of TCRs and LAT molecules, both of which can diffuse freely in the membrane (fig. 1B). Other signaling- and condensation-related molecules such as Lck, ZAP70, SOS, Grb2, PLCγ, and CD45 are modeled implicitly by the effects they have on either TCRs or LAT molecules. In each time-step of a simulated signaling trajectory, the actions of each TCR and LAT molecule are calculated independently, based on the probabilities that each molecule performs any of the available actions. The determined probabilities are then used to stochastically make the relevant changes to the state of the simulation grid, and the next time step begins. By explicitly modeling the location of each molecule our model can capture signaling effects that occur due to spatial inhomogeneities that are not considered in existing models.

**Figure 1.**
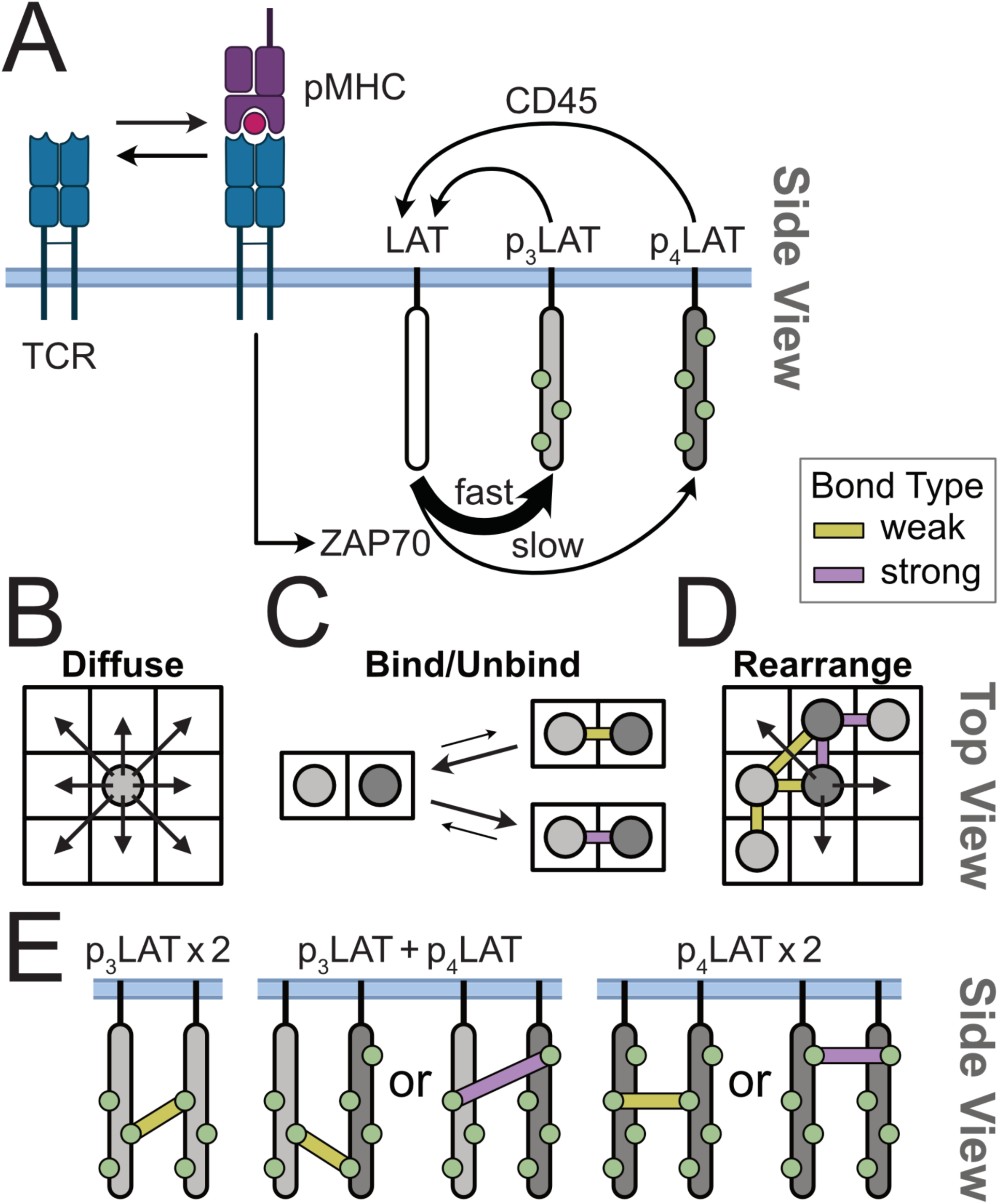
pLAT Condensation Model. A) The TCR module of the model consists of pMHC binding and subsequent activation of ZAP70. Active ZAP70 phosphorylates LAT to produce p_3_LAT (light gray) and p_4_LAT (dark gray). pLAT can be dephosphorylated by phosphatases such as CD45. B) The simulation takes place on a 2D grid over which pLAT molecules can diffuse. C) Adjacent pLAT molecules can bind to each other via cross-linking molecules that are not explicitly modeled. Bonds can be weak (yellow bar; slow on-rate, fast off-rate) or strong (purple bar; fast on-rate, slow off-rate). D) Molecules within a pLAT cluster can rearrange internally as long as the number of bonds made between them is conserved. E) Strong bonds (purple bars) require that at least one of the bonded pLAT molecules be p_4_LAT, while weak bonds (yellow bars) do not have this requirement.

To interrogate the role of LAT condensation independently from other signaling steps, we do not explicitly model the phosphorylation steps between TCR binding and ZAP70 activation. Instead, we assume that a pMHC-bound TCR immediately phosphorylates LAT (via ZAP70) in its local neighborhood, assuming that TCR phosphorylation and subsequent ZAP70 recruitment occur rapidly compared to LAT phosphorylation (fig. 1A). This assumption is based on findings that LAT phosphorylation is a rate-limiting step for T cell signaling initiation^27,28^. In reality, upstream events do take time to occur, and are likely also involved in kinetic proofreading^29,30^, especially in suppression of signals from very short binding ligands (e.g. binding dwell times less than a second). However, because we sought to evaluate the degree to which LAT condensation can contribute to kinetic proofreading and other signaling properties, we chose to leave out the effects of other phosphorylation steps on these properties.

We assume that the pMHC-bound TCR complex phosphorylates LAT at four pY sites^31^: three sites are ideal substrates for the kinase and hence are phosphorylated at a higher rate, while one (Y132) is less ideal and is phosphorylated at a lower rate^27,32^ (fig. 1A). We model the Y132 site separately from the other sites, as its slower phosphorylation has been shown to be a rate limiting step in T-cell signal initiation^27,28^. To account for these differences, we model two types of pLAT: p_4_LAT, which has all pY sites phosphorylated and can participate in PLCγ-mediated bonds, and p_3_LAT, which has no phosphorylation on Y132 and can bind other p_3_LAT only via weaker Grb2-SOS-mediated interactions (fig. 1A,C,E). These reversible interactions are multivalent and allow LAT to form a condensate by creating a network of bonds through its pY sites^10,20,33^.

pLAT condensate formation is opposed by the activity of phosphatases (such as CD45) which dephosphorylate pLAT, rendering it incapable of crosslinking (fig. 1A)^34,35^. While phosphatases act rapidly at uncondensed LAT sites, they are sterically hindered both directly by the binding of other molecules to pY sites, and indirectly through exclusion from condensates^20,21,36^. We thus modeled a slower dephosphorylation rate for pLAT that has more bonds to other pLAT, with the stronger binding of PLCγ to pY132 offering a higher degree of protection than the weaker Grb2/SOS mediated bonds.

Altogether, our model captures the key physical and chemical processes underlying TCR-induced LAT clustering, thus allowing us to assess how this emergent phenomenon contributes to the observed input/output characteristics of the TCR signaling pathway.

### Phosphorylated LAT condenses above a critical concentration

*In vitro* studies have shown that pLAT can undergo phase separation above a critical concentration^20^. We first ran simulations where the number of pLAT molecules was held constant by removing the ZAP70 and CD45 activity from the model to determine whether our model could recapitulate similar pLAT condensation behavior. In these simulations we expect to observe behaviors that resemble a phase transition, including the presence of a critical concentration below which clusters do not form, and above which they form rapidly^37,38^. Consistent with these expectations, we observed a sharp transition where pLAT began to form clusters above a critical concentration threshold (160/μm^2^; fig. S1A,B,C; movie S1,2). Additionally, at this critical concentration, pLAT shows a very clear bimodal distribution where each simulation either achieves complete clustering or does not form clusters at all, indicating that once a condensate is nucleated it forms very quickly and is very stable (fig. S1D). Finally, we found that the mean time to first cluster decreases with concentration following a power law with an exponent of roughly 3.5, suggesting that cluster nucleation requires simultaneous binding of 3-4 pLAT molecules (fig. S1E). These results demonstrate that the model recapitulates the key aspects of nucleation and condensation as intended and allows us to further explore how these features can contribute to the behaviors of the TCR signaling pathway as a whole.

### Individual bound TCRs nucleate signaling condensates with a time delay

Having shown that pLAT alone can undergo clustering in our model, we proceeded to ask whether pLAT can form clusters in response to TCR pMHC engagement and LAT phosphorylation. Single-molecule studies had previously shown that single TCRs can induce LAT clusters with a variable time delay after pMHC binding^24^. In these experiments TCRs are only tracked after they bind to a pMHC, and the peptides used are very high affinity. We therefore ran our simulations with the TCR constantly bound to pMHC throughout the simulation (fig, 2A,B) to mimic the tracked TCRs in the experiment.

From simulations, we found that single active TCRs can nucleate clusters of pLAT molecules with a time delay, consistent with experimental observations^24^ (fig. 2A; movie S3). In this representative simulation (fig. 2) we found that a small cloud of pLAT molecules formed under the active TCR very shortly after the start of the simulation, with a concentration gradient that decayed with increasing distance away from TCR. While pLAT molecules in this cloud could interact with one another, these interactions were transient, such that the majority of molecules were either monomers or dimers (fig. 2B, inset). However, after a time delay, a LAT cluster became nucleated and proceeded to grow, fed by continuing TCR activity (fig. 2B). Notably, these results show that a single pMHC ligand gave rise to a large cluster containing hundreds of LAT molecules. This remarkable amplification in the number of active signaling molecules is also observed experimentally^15,24^ and demonstrates the ability of a condensation-based signaling system to sensitively amplify signals from single molecule binding events.

**Figure 2.**
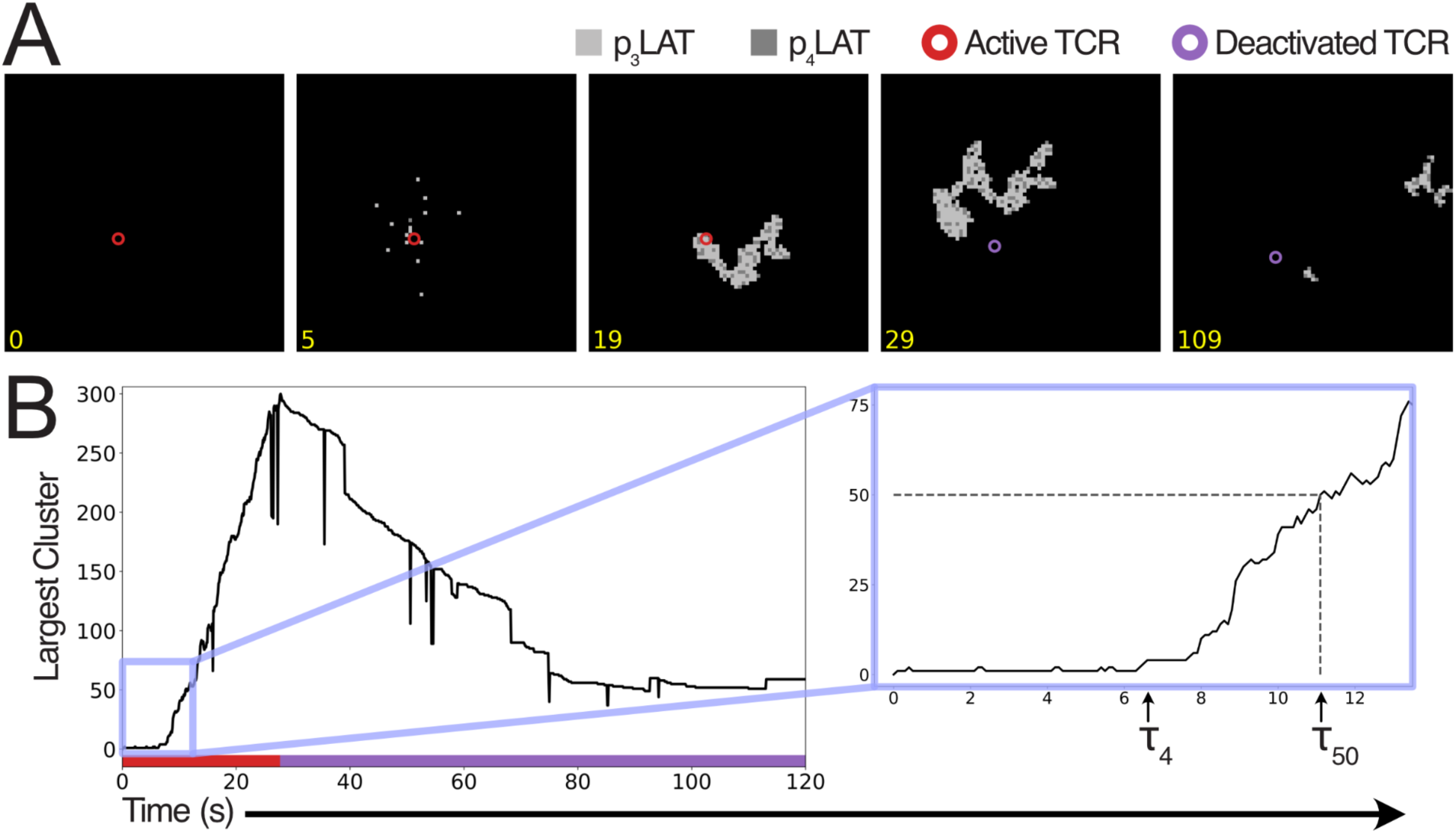
Individual bound TCRs nucleate signaling condensates with a time delay. A) Snapshots from an example simulation of a single permanently bound TCR. Light gray squares represent p_3_LAT and dark gray represent p_4_LAT. Circles represent TCRs, red when it is active, purple when it is deactivated. Each snapshot represents a 2.25μm square. Time stamps are in seconds. B) Trace of the size of the largest cluster over time in the example simulation in (A) showing the time to the first cluster of size 4 or greater (τ_4_), or size 50 or greater (τ_50_). Colored bar indicates TCR status; red for active, purple for inactive.

### Simulated LAT cluster sizes and dynamics match single-molecule imaging

To further validate our model, we compared LAT clustering dynamics from simulations to those previously seen in single molecule imaging experiments^24^ (fig. 3). These experiments track individual pMHC-TCR complexes and LAT clusters that form nearby; importantly, because they track the initial time that a single TCR binds to a pMHC ligand, they allow delay times and size distributions of pLAT clusters to be precisely measured. In order to compare cluster sizes between simulation and experiment, we define N_max_ to be the maximal number of pLAT molecules in clusters over the course of a simulation (fig. 3A). Our simulations show a range of N_max_ values from 50 to more than 500 molecules (fig. 3B), demonstrating the ability to amplify a single-molecule input signal into a much larger response. This range of sizes closely matches the distribution observed experimentally, where cluster sizes range from the detection limit of 200 molecules to about 700, with the majority between 250 and 400 molecules^24^.

**Figure 3.**
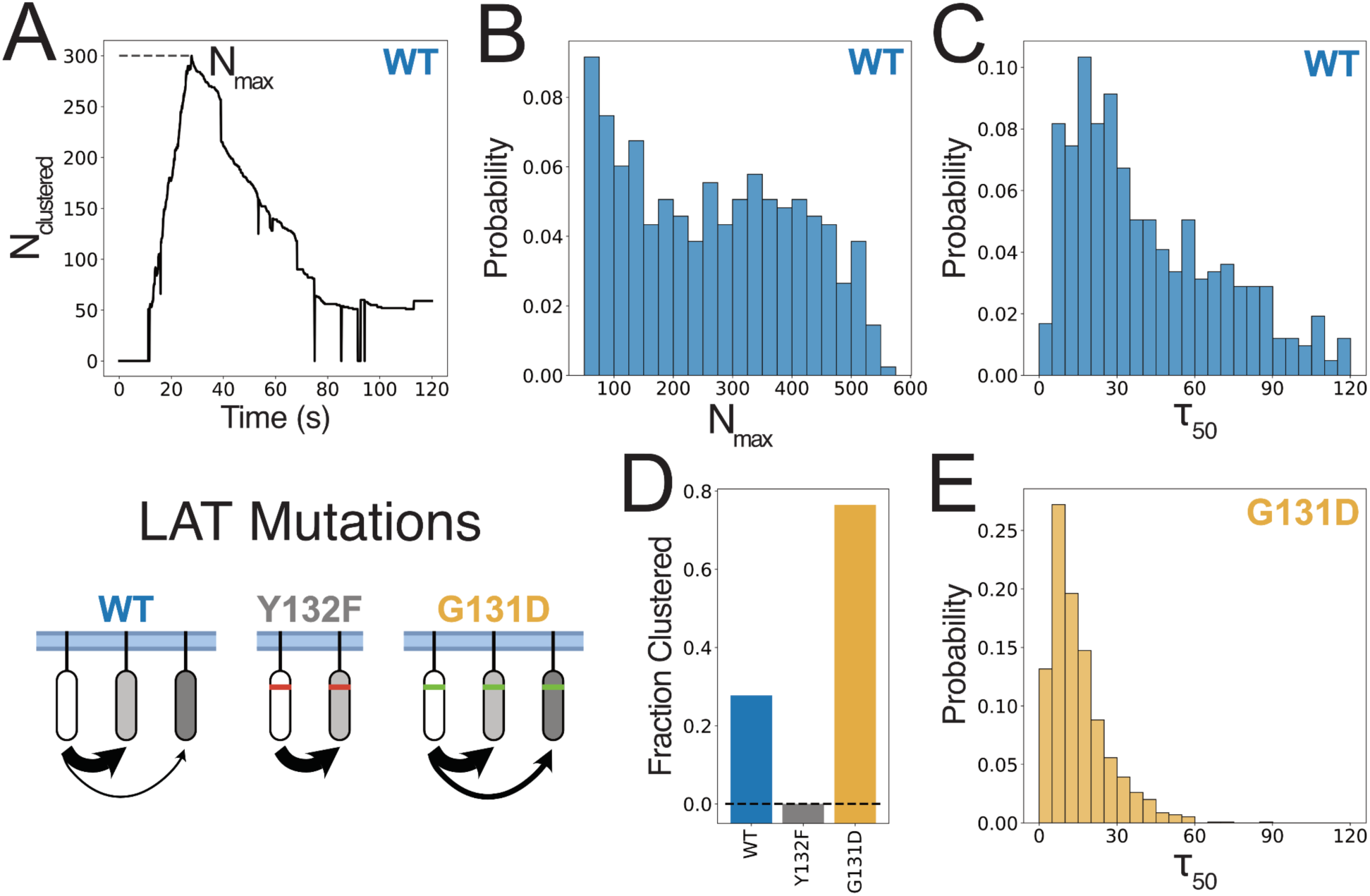
Simulated LAT clustering dynamics match experimental results. Data from simulations of single permanently bound TCRs on cells with varying LAT mutations. 1500 replicate simulations were run for each mutant. A) Representative trace of the number of pLAT molecules in clusters of size 50 or greater in a single simulation with WT LAT. N_max_ denotes the maximum number of clustered molecules over the course of the simulation. B) Distribution of N_max_ values over replicate simulations with WT LAT. C,E) Distribution of τ_50_ values (the time it takes to form a cluster of size 50 or greater) in simulations for either WT LAT (C) or G131D LAT (E). Simulations in which no cluster forms are omitted from these distributions. D) Bar chart showing the fraction of simulations that produce a LAT cluster for various LAT mutants.

We next quantified the delay time between pMHC binding and pLAT cluster formation (τ_50_, fig. 3C), as this time delay in signaling condensate nucleation could constitute a kinetic proofreading step in the TCR signaling pathway. Comparing the simulated and experimental distributions, we find that they are strikingly similar with an initial rise peaking around 20-25s, followed by a slow decay (fig. 3C)^24^. Critically, both distributions do not follow a monotonic decay such as the exponential distribution expected from a first-order process. This delay in LAT cluster nucleation could contribute to selectivity by filtering out short-lived binding events from weak ligands, a possibility we will test below.

Finally, in single-cell molecule experiments, pMHC-bound TCRs give rise to pLAT clusters only in a fraction of cases (25-50%), even when they were bound for very long times^24^. This cap on TCR-induced clustering likely arises from feedback mechanisms which deactivate pMHC-bound TRCs over time. As the binding lifetime increases, if a cluster has not yet formed, it becomes more likely that the TCR will be deactivated before a cluster can form. This negative feedback is incorporated into our model and sets the probability of clustering at about 27% (fig. 3D, WT), in accordance with experimental data^24^.

The probability of forming a cluster, the size of the cluster that is formed, and the delay between pMHC-TCRC binding and cluster formation are all critical parameters controlling the T-cell response at a cellular and population level. The close agreement in these measurements between our simulated results and experimental data suggest that our model is able to accurately capture these key aspects of early TCR signaling.

### LAT phosphorylation kinetics modulate the time delay to condensate nucleation

LAT Y132 phosphorylation has been shown to be a rate-limiting step in T-cell signaling initiation^27^, and LAT mutations that accelerate this reaction have further been shown to promote LAT cluster formation^24^. Here, we test whether changes to rates of Y132 phosphorylation can modulate time delays to condensate nucleation. To do so, we simulated two mutations in LAT which are known to affect both the PLCγ binding site and the kinetics of signaling. The glycine residue at position 131 in WT LAT makes Y132 a poor substrate for ZAP70, reducing its phosphorylation rate significantly^27^. The G131D mutation increases this rate^27^, which we modeled as an increase in the p_4_LAT production rate in our simulations. Conversely, the Y132F mutation completely prevents phosphorylation at the same site, modeled as a reduction of the p_4_LAT production rate to zero. We first observed that these mutations had the expected effect on the fraction of TCR binding events that are productive. Y132F completely abolishes clustering in simulations (fig. 3D), paralleling the dramatic reduction in signaling observed in T-cells bearing that mutation^31,39^. Conversely, G131D increases the productive binding rate from 27% to about 75% (fig. 3D), in agreement with the observation that T-cells bearing this mutation are more easily activated^27^.

In addition to affecting the probability of clustering, G131D dramatically reduces nucleation delay times. Both simulated and experimental results show an earlier peak in the delay time distribution around 15s and a shorter tail to the right of the distribution (fig. 3E)^24^. The shortened delay suggests a decreased ability for G131D cells to discriminate between weak and strong binding pMHCs.

Through these simulations we show that our model recapitulates the effects of mutations in LAT that affect the kinetics of its phosphorylation at the key Y132 site. We are able to recapitulate expected effects on the frequency of cluster formation as well as on the distribution of cluster initiation times. And more broadly, we are able to show that the kinetics of LAT phosphorylation have a direct influence on the signaling outcome, and are likely to influence kinetic proofreading via modulation of delay time distributions.

### Kinetic proofreading by condensate nucleation in TCR signaling

The observed delay between pMHC binding and LAT condensation can potentially serve as a kinetic proofreading step and thereby enable T-cells to selectively respond to high-affinity pMHC ligands. To test this idea, we simulated a system with diffusible TCRs on a 2D membrane that were then stimulated with pMHCs with varying lifetimes of TCR binding. In order to allow a fair comparison between different affinity pMHCs, we increased the simulated concentration of pMHC inversely with its TCR binding lifetime, thus keeping TCR occupancy the same (fig. 4A,C). In these simulations, the fraction of pMHC-bound TCRs remained the same as binding lifetimes increased, as expected (fig. 4C). In contrast, LAT clustering occurred preferentially in response to stimulation with tight-binding pMHCs, with little or no clustering occuring for pMHC ligands with shorter binding lifetimes (τ_off_ < 1s) (fig. B, D-E; movie S4-10). Furthermore, as pMHC binding lifetimes increased, both the likelihood of cluster formation as well as cluster sizes increased, indicating that the system is able to encode pMHC affinity information at this step. These results directly demonstrate kinetic proofreading by LAT condensation. Because the TCR occupancies are matched across all simulations, the reduced clustering observed with the faster off-rates can only be due to a kinetic proofreading mechanism. To the extent that this occurs in these simulations, the proofreading is carried out solely by LAT condensation, and not any upstream components of the pathway since they are not considered in our model. These results show that LAT condensate nucleation is sufficient to generate the time delays needed to enable selective response to only high-affinity pMHC ligands.

**Figure 4.**
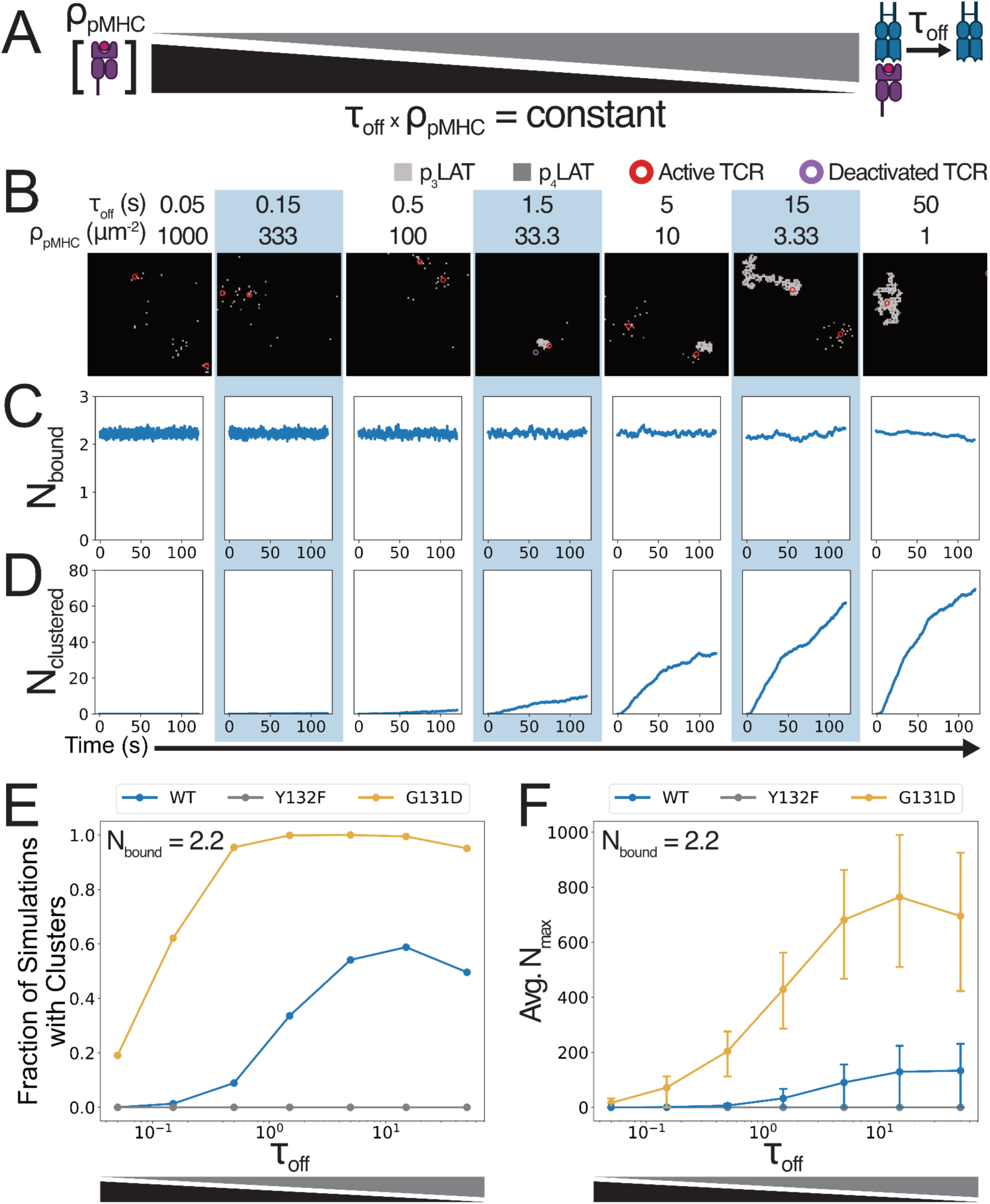
Kinetic proofreading by condensate nucleation in TCR signaling. Data from simulations for varying LAT mutants and pMHC lifetimes, with TCR occupancy held constant. 750 replicate simulations were run for each condition. A) Schematic showing the simultaneous increase in pMHC-TCR bond lifetime and decrease of pMHC 2D density used to achieve constant numbers of bound TCRs. B) Snapshots of representative simulations at each pMHC off-rate with WT LAT. Light gray squares represent p_3_LAT and dark gray represent p_4_LAT. Circles represent bound TCRs, red for active and purple for deactivated. Each snapshot represents a 2.25μm square. C) Average number of bound TCRs over time for simulations at each pMHC off-rate with WT LAT. D) Average number of pLAT molecules in clusters of size 50 or greater for the same simulations in (C). E) The fraction of simulations for each condition that produce a cluster of size 50 or greater. F) Mean N_max_ over replicate simulations for each of the simulated conditions. Error bars show 25^th^ and 75^th^ percentiles of replicate simulations.

To probe the role of slow LAT phosphorylation in setting time delays in kinetic proofreading, we ran the same TCR occupancy-matched simulations with the G131D and Y132F mutants and looked at the effects of these mutations on signaling magnitude and proofreading. As expected, the G131D mutation increased the LAT clustering response both in terms of the fraction of simulations that were able to form clusters (fig. 4E) and the size of those clusters (fig. 4F), while the Y132F again completely prevented clustering. Strikingly, the G131D mutation drastically reduces the ability of the simulations to distinguish self-peptides (τ_off_<1s) from foreign peptides (τ_off_>10s) with roughly 20% of G131D simulations responding to peptides with 50ms half-lives (fig. 4E). The reduced degree of proofreading caused by this mutation explains why mice bearing this mutation show enhanced responses to weak pMHC stimuli and develop autoimmune disease^30^.

Together, these results indicate that LAT nucleation constitutes a key kinetic proofreading step in T cell signaling. The delay imposed by the rarity of nucleation events provides a simple mechanism for filtering out short-lived pMHC-TCR interactions. Additionally, the phosphorylation rate of the PLCγ binding site is a critical parameter that controls the threshold for binding lifetimes that are able to produce a condensation response.

### Cluster growth allows wide dynamic range

In addition to distinguishing between ligands of varying affinity, T-cells also need to respond to variations in pMHC levels^8^, which may indicate the severity of the infection. Most current models propose that while signaling is digital at the single cell level, the population can have an analog response where the pMHC dose tunes the percentage of cells that respond^4^. However, some aspects of TCR signaling have been demonstrated to respond to pMHC dose in a graded manner at the single cell level^8,40^.

To determine the extent to which LAT clustering can convey information about pMHC dose, we tested the response of our model to antigen doses spanning three orders of magnitude (fig. 5). We first found that each simulated cell has a binary response in terms of the presence or absence of a cluster, with the fraction of responding cells increasing as with the antigen level and lifetime (fig. 5A). At low doses, the average size of clusters, when they are present, remains low, but as the fraction of cells responding begins to saturate, the size of the cluster begins to grow (fig. 5B). These results indicate that our model is able to produce both all-or-none signaling responses in the nucleation of a LAT cluster, as well as graded increases in the sizes of nucleated LAT clusters with increasing antigen dose. Together, these results explain how LAT clustering dynamics could give rise to digital activation for signal amplification and single molecule sensitivity, as well as analog tuning for dynamic range in pMHC dose detection.

**Figure 5.**
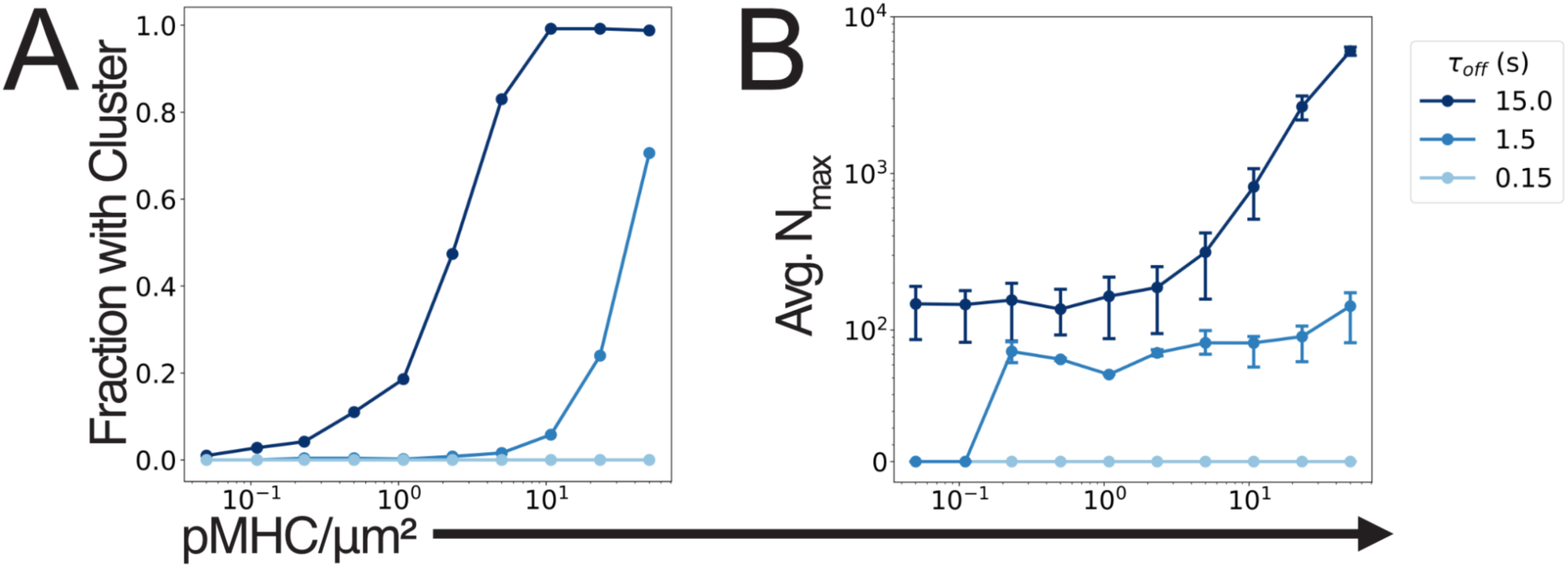
pLAT condensation increases gradually with increasing pMHC stimulation. Data from simulations with varying pMHC dose and lifetime, with WT LAT. 500 replicate simulations were run per condition. A) The fraction of simulations that produce pLAT clusters in response to stimulation with varied pMHC density at each of three TCR-pMHC half-lives. B) The mean N_max_ in simulations where a cluster forms for the same stimulation conditions as (A). Error bars show 25^th^ and 75^th^ percentiles of replicate simulations.

### LAT condensation sets an optimal pMHC spacing for T cell activation

The analysis above shows that our LAT condensate nucleation model can uniquely give rise to sensitivity, specificity and dynamic range in TCR signaling. This model agrees with existing experimental data; however, we have not yet shown that there are unique predictions made by this model that can allow it to be distinguished from other existing kinetic proofreading models. Most other dynamic models make the assumption of a well-mixed system, where a scalar value can represent the concentration of each signaling molecule. In these models, the number of long-lived binding events is the only relevant parameter for activation^4,7,12,13,15^, and parameters such as the spacing between antigens would not affect signaling activity, despite evidence that this may be important^41–43^.

To test if our model made any unique predictions about spatial dependencies in TCR signaling, we ran simulations where a square grid containing a fixed number of pMHCs was placed opposing the T-cell so that TCRs could only bind and activate in those locations (fig. 6A). We varied the 2D density of pMHCs over roughly three orders of magnitude by adjusting the spacing between them, while keeping the number of pMHCs constant, and examined the effects on LAT clustering (fig. 6B-C). In contrast to other models, our simulations show that there is an intermediate spacing of pMHCs that is optimal for LAT cluster formation and T-cell signaling (fig. 6C). At both low and high pMHC density the cluster sizes are smaller, while only intermediate densities achieve maximal clustering (fig. 6C; movie S11-15). At very low pMHC densities, each pMHC-TCR complex is isolated from others such that pLAT produced at one TCR is dephosphorylated before it can reach a cluster formed at another TCR (fig. 6D. As the pMHC density increases, TCRs are able to cooperate, adding pLAT to clusters that were nucleated by other TCRs, allowing the cluster to grow larger than what could be produced by a single pMHC-TCR complex (fig. 6E). However, if the pMHCs are brought too close together, then the TCRs become crowded, merging into what is effectively a single TCR that produces pLAT at a higher rate. This allows a cluster to form faster, but the cluster size is still limited to what can be produced by a single TCR (fig. 6F). Thus, LAT condensation sets the spatial scales of TCR activation in addition to the temporal scales discussed earlier.

**Figure 6.**
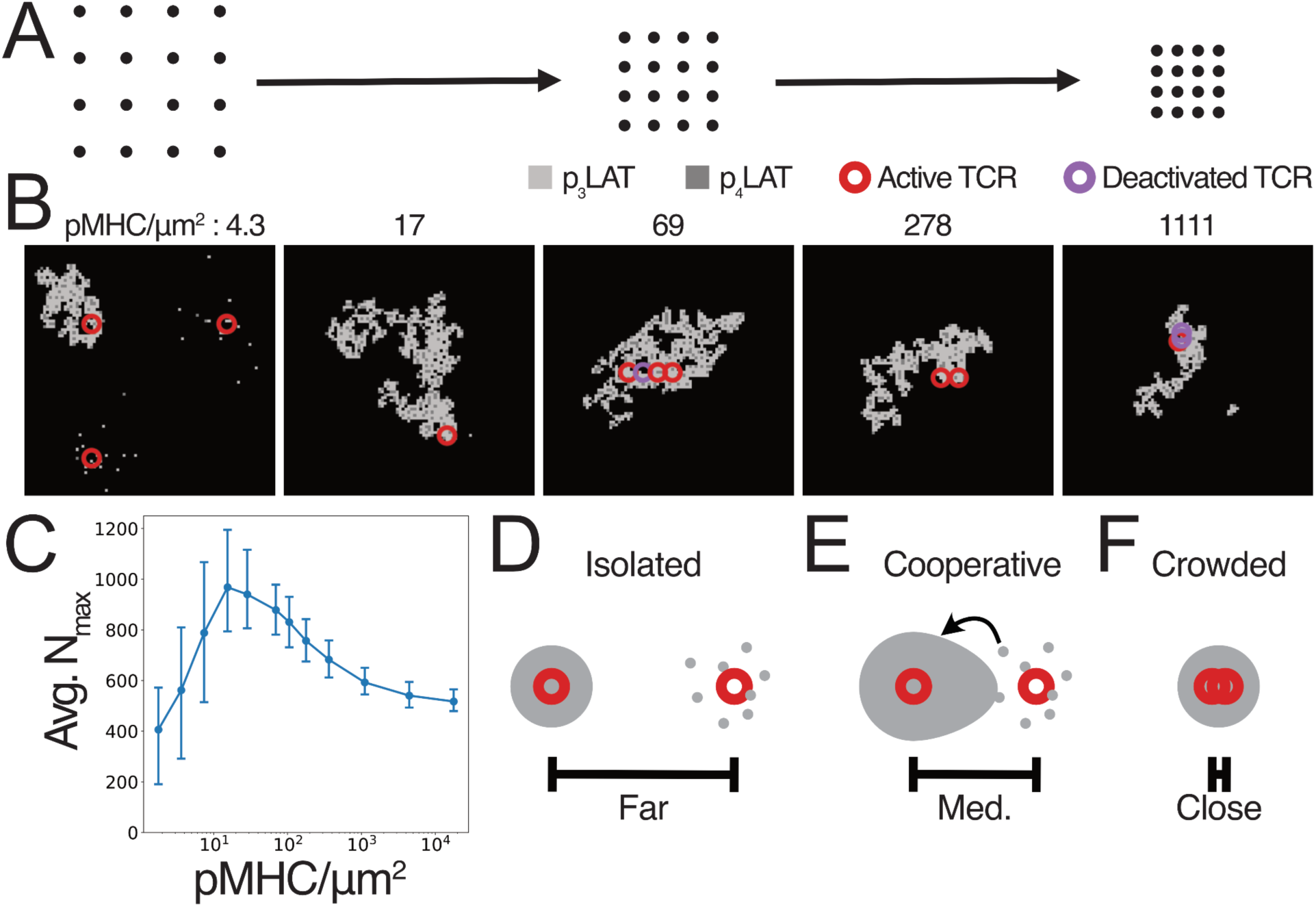
LAT condensation simulations predict an optimal pMHC spacing for TCR activation. Data from simulations with 16 fixed pMHCs at varying spacing, with WT LAT. 500 replicate simulations were run for each pMHC density. A) Schematic of pMHC placement in simulations of varying antigen density at constant antigen number. B) Snapshots from example simulations at different pMHC densities. Light gray squares represent p_3_LAT and dark gray represent p_4_LAT. Circles represent bound TCRs, red for active and purple for deactivated. Each snapshot represents a 1.27μm square. C) Mean N_max_ values for each pMHC density. Error bars show 25^th^ and 75^th^ percentiles of replicate simulations. D-F) Cartoon representation of the underlying cause of optimal spacing. Large gray circles represent pLAT clusters, small gray circles represent individual pLAT molecules, and red circles represent TCRs. D) At low pMHC density, pMHC-bound TCRs are isolated, preventing pLAT produced at one TCR from augmenting a condensate at the neighboring TCR. E) At intermediate pMHC density pLAT produced at one pMHC-bound TCR can add to a condensate nucleated at a neighboring TCR. F) At high pMHC density, neighboring pMHC-bound TCRs are crowded, limiting the area over which a condensate can form.

### 2D protein arrays allow precise tuning of pMHC spacing independent of dose

In order to test the prediction of optimal pMHC density, we developed an imaging assay to probe the relationship between antigen density and LAT clustering. As a platform for antigen presentation we used a 2D protein array^26^ to place pMHCs at regular intervals (fig. 7A). The arrays consist of two components in addition to the pMHC: the A component which is fused to the SpyCatcher protein^44^, and the B component which is fused to GFP. When mixed, the A and B components self-assemble into a hexagonal grid (fig. 7A). By controlling the stoichiometry of the SpyTag-pMHC and SpyCatcher-A prior to assembly with component B, we can control the 2D density of pMHCs on the arrays. Furthermore, by quantifying stochastic variations in the sizes of arrays we are able to assess effects of variations in pMHC number, thus allowing us to independently assess the effects of pMHC spacing and number on TCR signaling.

**Figure 7.**
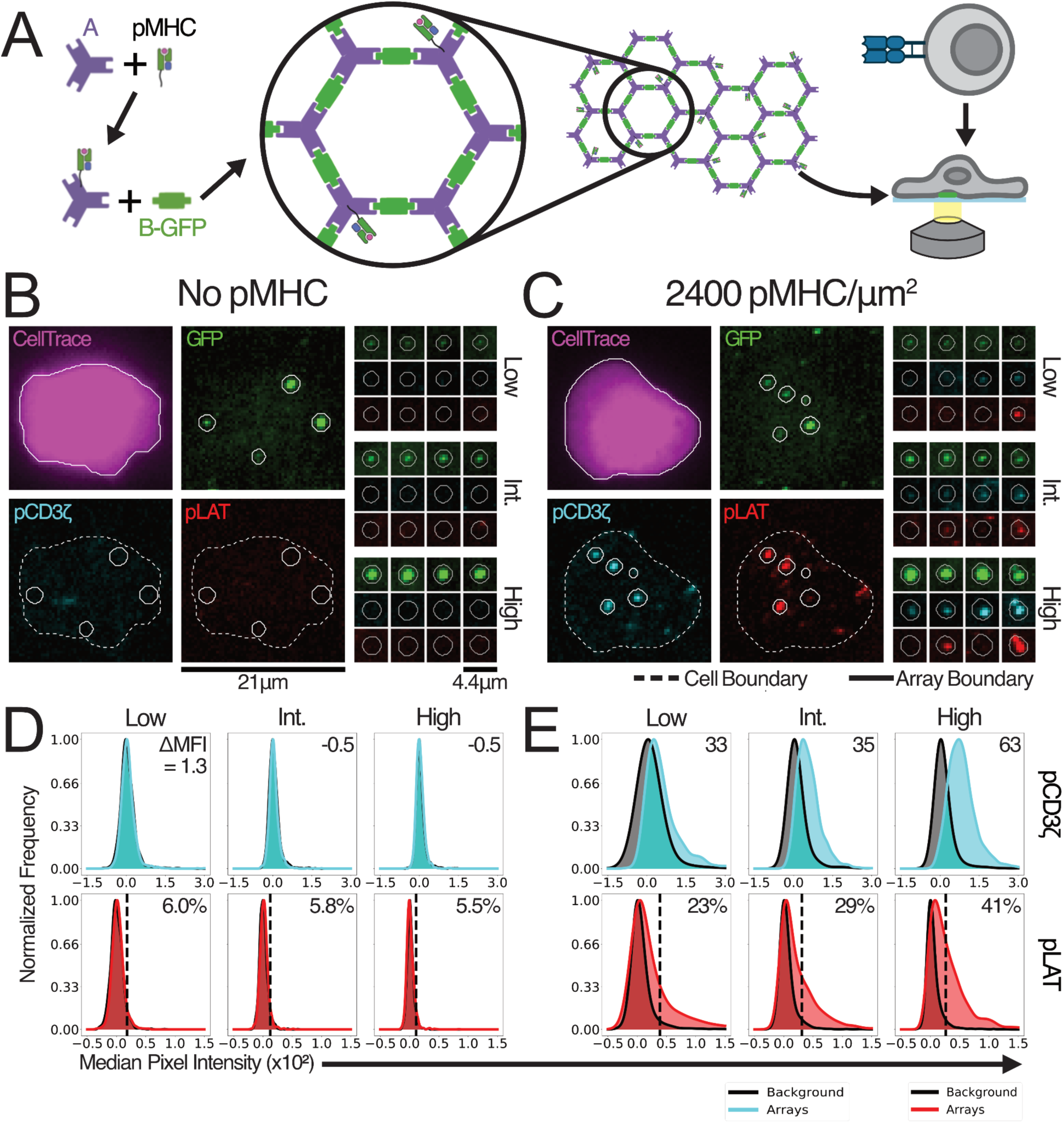
2D protein arrays enable precise pMHC display. A) Illustration of the array assembly and imagining protocol. Components A and B are shown as purple triangles and green rectangles, respectively. The assembled array has a 31nm distance between opposing edges of each hexagonal unit^26^. B,C) Example images of cells (purple) and arrays (green) that were fixed 5min after interacting with blank (B) or pMHC-presenting arrays (C) showing the pCD3ζ (light blue) or pLAT (red) responses to pMHC array stimulation. Larger fields of view (left) show a single representative cell interacting with arrays. Smaller fields of view (right) show four representative arrays with either low (33^rd^ percentile), intermediate (int.; 66^th^ percentile), or high (100^th^ percentile) GFP intensities. D,E) Normalized kernel density estimates for the distributions of median pCD3ζ (light blue, top) or pLAT (red, bottom) signal for arrays with low (left; 0-33^rd^ percentile) intermediate (middle; 33^rd^-66^th^ percentile) or high (right; 66^th^-100^th^ percentile) GFP intensities, compared to the same measurement in random locations (gray) for blank (D) or pMHC-presenting arrays (E).

As a model antigen, we presented the gp33-41 peptide from Lymphocytic Choriomeningitis Virus (LCMV) complexed to the murine class I MHC, H-2D(b)^45^. We used as a model a Jurkat T-cell line expressing the cognate P14 transgenic TCR and the CD8 co-receptor, which we found gives rise to robust downstream signaling^8^. After the arrays were assembled, we seeded them onto a glass imaging surface and plated the P14 Jurkat cells on top of them. After a brief stimulation, we fixed the cells and stained them for phospho-CD3ζ (pCD3ζ) and phospho-LAT (pLAT; fig. 7B,C). Each array produced a bright GFP spot in the image which we used to locate the regions of the image where arrays overlapped cells. We additionally calibrated our measurements of GFP brightness in order to be able to count the number of GFP molecules and, by extension, the number of pMHCs present in each array (fig. S2)^46^. Using these values, along with several other measurements, we filtered the arrays to remove aggregates and imaging noise, yielding a set of high quality arrays that we used in all downstream analysis (fig. S3).

We first used the median phosphorylation signal in each array as a measure of the effectiveness of array stimulation (fig. 7D,E). These measurements showed a significant elevation of pCD3ζ and pLAT levels in arrays that have a high density of pMHC on them, while those with no pMHC did not induce any phosphorylation above background (fig. 7D,E). Additionally, larger arrays (higher GFP copy number) induced higher levels of phosphorylation (fig. 7E). These results confirm that we were able to stimulate Jurkats with the arrays in a pMHC-dependent manner, and that phosphorylation is dependent on the number of pMHCs that contacted the cell. We note that array-displayed pMHCs differ from those displayed on cell membranes in their TCR contact heights, as well as their ability to exclude membrane-associated proteins such as phosphatases. These kinds of differences have been shown to impact TCR signaling initiation^42,47^ and could affect the fit of the model to the resulting data because the model was adjusted to match data from membrane-bound pMHC stimulation. We therefore compare only general trends in these experiments with the model predictions instead of precise values.

Consistent with the differing biochemical mechanisms for activation of CD3ζ and LAT, the distributions of their phosphorylation levels show different shapes in response to pMHC stimulation (fig. 7E). The pCD3ζ distribution shows a clear increase in its mode relative to the background distribution as the array size increases, consistent with CD3ζ phosphorylation occurring in proportion to the number of pMHCs present. The pLAT distribution, on the other hand, shows a widening tail to the right without a major shift of the mode. This distribution is consistent with pLAT signals resulting from stochastic cluster nucleation in a fraction of cases that increases with pMHC number. These results are consistent with our model prediction that LAT signaling condensates nucleate only for a fraction of pMHCs (fig. 3C), but can grow to large sizes that scale with the number of initiating TCR inputs (fig. 5).

### pMHC arrays reveal optimal antigen spacing for LAT condensation

With the functionality of the 2D pMHC arrays validated, we moved forward to test whether varying pMHC density has an impact on downstream signaling. To do this we created arrays with pMHC density varying from 37.5 to 2400 pMHC/μm^2^ and measured the CD3ζ and LAT phosphorylation responses to each density. We first examined the relationship between array size and phosphorylation signal and found that, for all pMHC densities above zero, phosphorylation increased with increasing array size (fig. S4A). The slope of this relationship also increased with increasing pMHC density (fig. S4B). These results show increasing pMHC number results in a stronger downstream CD3ζ and LAT phosphorylation; more antigen results in a stronger signal.

After demonstrating that pMHC array stimulation produces the expected response to varying pMHC number, we next tested whether TCR signaling varies with pMHC spacing at fixed pMHC number. To compare arrays across all pMHC densities despite non-overlapping distributions of pMHC number when comparing the lowest densities to the highest, we measured the activation efficiency for arrays of each density. This metric is defined as the slope of the line of best fit to the relationship between phosphorylation intensity and pMHC number (fig. S5), and measures the degree to which pMHCs elicit signaling, at a given spacing. We observed that pMHC spacing does not have a strong effect on CD3ζ activation efficiency (fig. 8A), while pLAT activation efficiency shows a clear peak at intermediate pMHC spacing (fig. 8B). This behavior is very similar to the behavior predicted by our model (fig. 6). The phosphorylation of CD3ζ, which does not participate in LAT condensation, is not impacted by pMHC density. Conversely, LAT phosphorylation is dependent on pMHC density, and requires an optimal density for maximal signaling. These experimental and simulated results further demonstrate that LAT condensation is central to the TCR signaling pathway. By examining the experimental and simulated responses to varied antigen density we have demonstrated that the optimal antigen spacing is controlled by the underlying length scales of LAT clustering. Thus, LAT condensation controls both the length and time scales required for optimal T-cell activation.

**Figure 8.**
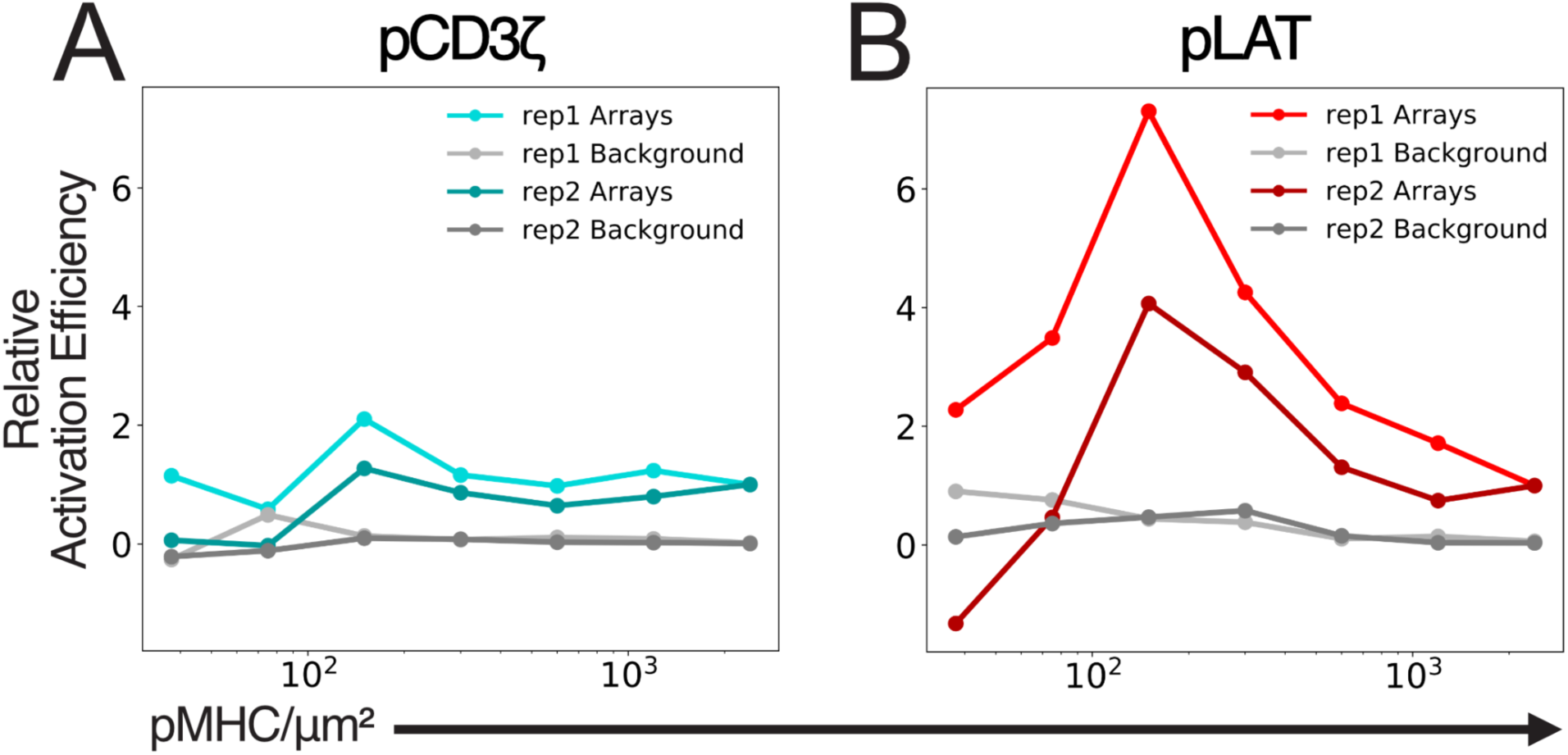
2D arrays reveal optimal pMHC spacing for LAT phosphorylation. A,B) Line plots showing CD3ζ (A) or LAT (B) activation efficiency as a function of pMHC density for arrays (color) or background (gray). Activation efficiency is defined as the slope of the line of best fit for the relationship between integrated phosphorylation intensity and pMHC number. Data from two biological replicates (rep1 and rep2) are shown, with each replicate independently normalized to the efficiency at 2400pMHC/μm^2^.

## Discussion

The results presented here demonstrate that condensation can explain the key features of the TCR signaling network, simultaneously achieving sensitivity, selectivity and dynamic range. Prior work in modeling these properties has generally focused on the selectivity of the pathway^4,7,12,13,15,19^ with some studies also giving attention to the sensitivity^4,13,16^ and very few considering dynamic range^7^. The focus on selectivity likely stems from the key role it plays in preventing autoimmunity, and the fact that it is difficult to balance all three of these properties in a single model. Each property imposes its own set of constraints on the underlying signaling network, and these constraints are often in opposition to each other. Large numbers of proofreading steps have been shown to increase the selectivity of the pathway^12,15^, but recent work has demonstrated that these additional steps make the system much more noisy, and can actually decrease the ability of the pathway to distinguish between weak and strong binding amid the stochasticity of biochemical reactions^19^. Strong positive feedback can significantly improve the sensitivity of a signaling pathway, but can create a digital system that is either “on” or “off” and lacks dynamic range. Strong positive feedback is also more likely to amplify small, spurious responses to off-target signals, reducing specificity. Conversely, negative feedback can improve selectivity and can explain the observation that antigens with intermediate affinity can antagonize the signaling of higher affinity antigens^4,13^. The ability of these additional feedback loops to strengthen our model’s proofreading and improve its performance could be explored in future work.

Our model is able to balance the constraints imposed by each property of the TCR signaling pathway in a single signaling step by taking advantage of the inherent physical properties of condensation. Simulations of single pMHC-TCR interactions show how condensation itself provides a strong positive feedback mechanism that can amplify even a single pMHC binding event into a large LAT cluster. The resulting cluster is able to grow in proportion to the amount of available pLAT, giving the system a wide dynamic range. We have also demonstrated that the positive feedback inherent in condensation is carefully controlled by another condensation-related mechanism: nucleation delays. Because of strong ambient phosphatase activity, pLAT is very short-lived outside of clusters resulting in fast dissipation of any pLAT that accumulates during short pMHC binding events, preventing spurious activation and filtering out noise. However, if a pMHC can bind long enough that a LAT cluster can nucleate, that cluster can grow quickly and survive even after the pMHC dissociates, allowing time for downstream signaling to be fully activated. In addition to these well-recognized properties of the TCR pathway, we have demonstrated that the same condensation mechanism underlies the spatial preferences of TCR signaling. The area over which a single TCR can produce a LAT cluster sets the optimal distance between active TCRs. The preference for intermediate antigen spacings that we observed in our model and imaging experiments is also consistent with previous work which used DNA origami to cluster defined numbers of pMHCs together^43^. The DNA origami results showed that clusters with intermediate pMHC numbers are most efficient for downstream activation^43^.

It is striking that a single step in this complex signaling pathway can achieve such a wide variety of signaling properties in contrast to other models that require a complex multi-step network of reactions to achieve the same results^4,7,12,13,15^. Our model provides an intuitive explanation for the properties of TCR signaling that make T-cells such an effective component of the immune system. It also provides insight into how the system is carefully tuned to its purpose, and how small adjustments to key parameters can lead to dysfunction. The insights provided here not only paint a more complete picture of the TCR signaling mechanism, but also provide avenues to adjust T-cell signaling in therapeutic contexts. By carefully controlling LAT clustering, T-cell signaling could be modulated to suit the situation. In this way, our model can be used to both further the understanding of TCR signaling and provide guidance on how to best engineer it to fill therapeutic needs.

## Methods

### Computational model

Our computational model is based on a 2D square grid representing the inner leaflet of the plasma membrane. We chose to use a grid spacing of 30nm to approximately match the space a single molecule of LAT occupies in the densest LAT clusters observed experimentally. Within this grid, TCRs and pLAT molecules are modeled as occupying a single grid square to the exclusion of molecules of the same type. All other molecules are not present in the model, and their effects are modeled implicitly in the behavior of the TCRs and pLAT molecules. Each timestep of the model proceeds through the following steps:

1. TCR diffusion
2. pLAT dephosphorylation (CD45 activity)
3. TCR-pMHC binding and activation
4. LAT phosphorylation (ZAP70 activity)
5. pLAT diffusion
6. Intra-cluster rearrangement
7. pLAT-pLAT binding (via Grb2, SOS, and PLCγ)

These steps are repeated for the desired number of time steps (for a total of 2-5min of simulated time). The length of the time step is set to 125μs so that the probability that any molecule would take more than one action within each step of the simulation is negligible. Further details of each step are outlined below:

1. TCR Diffusion TCRs diffuse across the simulation grid from one square to an unoccupied adjacent or diagonal square with a probability, P_dif_,_f_ defined below:

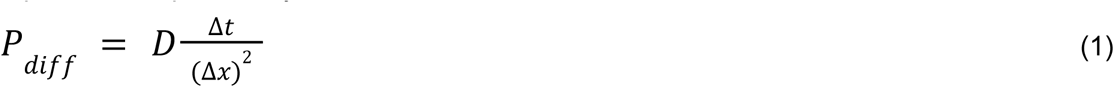

Where Δt is the simulation time step and Δx is the width of the grid square, and D is the diffusion coefficient of the TCR. Based on experimental measurements, we set the diffusion coefficient of pMHC-bound TCRs to be roughly 15 times slower than free TCRs^27^.
2. pLAT Dephosphorylation The activity of CD45 and other phosphatases is incorporated into the model as the dephosphorylation probability of pLAT. Since the pY sites on LAT are bound by other proteins in the cluster, the probability of dephosphorylation for an individual pLAT molecule is reduced as it forms more bonds, according to the following equation:

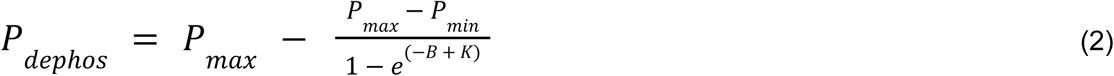

Where P_max_ and P_min_ are the bounds on the probability of dephosphorylation, K is a constant that determines how many bonds are required for protection, and B describes the number and type of bonds made by the molecule:

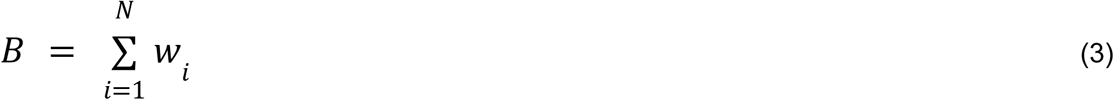

Where N is the number of bonds made by the molecule and w_i_ is the weight of the i^th^ bond. Bond weights are high for the strong interactions mediated by PLCγ and low for the weaker interactions mediated by Grb2 and SOS. The protection from phosphatases provided by a cluster is the basis for one of several positive feedback mechanisms in condensate formation.
3. TCR-pMHC binding and activation TCR binding occurs through standard binding kinetics with second order kinetics for association and first order kinetics for dissociation. In each simulation, only TCRs or MHCs, but not both, are explicitly modeled, and their binding probability is modulated by a parameter which represents the 2D density of their unmodeled binding partner. In addition to being bound or free, TCRs can exist in a signaling competent or incompetent state. All TCRs start signaling competent, meaning that they can immediately activate ZAP70 and begin phosphorylating LAT as soon as they bind a pMHC. However they become signaling incompetent with first order kinetics during the times that they are bound to a pMHC. This accounts for the local negative feedback that has been observed to occur on TCRs in single molecule imaging studies^24^.
4. LAT phosphorylation In order to focus on the contributions of LAT clustering to TCR signaling and simplify the model, active TCRs (pMHC-bound and signaling competent) are assumed to instantly phosphorylate ZAP70 which can then phosphorylate LAT. Each active TCR can phosphorylate LAT in the 3x3 grid centered on its location with a probability proportional to the number of sites in the 3x3 neighborhood that are not occupied by pLAT, representing the local substrate depletion around an active TCR. When pLAT is produced, it can be made in one of two forms representing different sites of phosphorylation. pY132 is modeled separately from the other three pY sites since it is known to behave differently. Primarily, pY132 is known to be phosphorylated more slowly by ZAP70^27,28^, so pLAT is more often produced with only the other three sites phosphorylated (p_3_LAT) while the fully phosphorylated form (p_4_LAT) is produced more rarely.
5. pLAT diffusion pLAT diffusion proceeds very similarly to TCR diffusion as described in step 1, but with a faster diffusion coefficient that is modulated by the number of molecules that move together in a cluster:

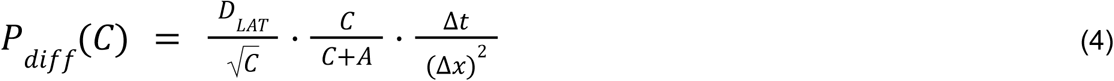

Where D_LAT_ is the diffusion coefficient for a single pLAT molecule, C is the number of molecules that are in the cluster, and A is the number of additional molecules that are pushed by the cluster as it moves. The first term models the reduced diffusion coefficient for larger objects embedded in the membrane. The second term accounts for the inertia of molecules not in the cluster as it collides with them. The third term accounts for the dimensions of the simulation as in equation 1.
6. Intra-cluster rearrangement Once in a cluster, pLAT molecules are allowed to rearrange within the cluster to model the flexibility that has been observed experimentally in LAT clusters^20,21^. This rearrangement happens at a much slower rate than free diffusion, and can only occur when the new location of the pLAT molecule allows it to make the same number of each type (PLCγ or Grb2/SOS) of bond with its new neighbors as it did before the rearrangement.
7. pLAT-pLAT binding Adjacent molecules of pLAT can bind to each other through the implicit activity of PLCγ, Grb2, and SOS. Any molecules that are connected directly or indirectly by a bond must move together as a cluster until the bonds connecting them are broken. These bonds can either be strong bonds mediated by PLCγ, or weaker bonds mediated by Grb2 and SOS. However, since PLCγ binds preferentially to pY132 on LAT, at least one p_4_LAT must participate in any strong bond. The strength of the bond is modeled by the first-order off-rate of the bond, with stronger bonds having slower off rates. In contrast, the on-rate of the bonds is modulated by their local environment to account for the local concentration of PLCγ, Grb2, and SOS. As a pLAT molecule forms more bonds, the local concentration of the crosslinking molecules is increased, thus increasing the effective on-rate of modeled pLAT-pLAT bonds. This increase on the on-rate drives a positive feedback loop which allows pLAT condensates to grow quickly once nucleated, following the equation below:

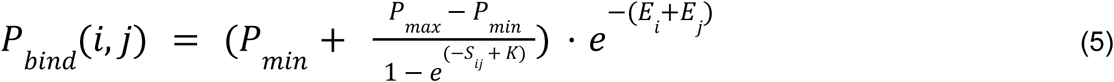

Where P_max_ and P_min_ are the bounds on the probability of binding, K describes the degree of local binding required to promote further binding, and E_i_ and S_ij_ are defined as follows: E_I_ is equal to 0 if molecule i is already bound to a molecule other than j, and is a positive number, Z, representing the entropic penalty of binding if molecule i is free.

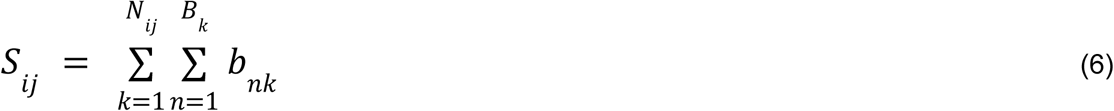

Where N_ij_ is the number of pLAT molecules that neighbor both molecules i and j, B_k_ is the number of bonds made by molecule k, and b_nk_ is the weight of the bond between molecules n and k. Thus, as more bonds are made around a pair of pLAT molecules, the local concentration of cross-linking molecules increases, resulting in an increased probability of binding. Because PLCγ-mediated bonds only require a single molecule to form, the P_min_ for the formation of those bonds is increased relative to the Grb2/SOS-mediated bonds.

### Array component purification

SpyCatcher-fused component A (A-SC) and GFP-fused cyclic component B (B-c-GFP) were expressed in *E. coli* as previously described^26^. *E. coli* cell pellets were resuspended in TBS supplemented with 5% (w/v) glycerol (TBSG), 1mM PMSF, and 0.7U/mL RNase A and lysed by sonication. Lysate was clarified by centrifugation at 14000xg for 30min and His-tagged array components were purified form the soluble fraction by Ni affinity chromatography in TBSG with either 40mM imidazole (wash) or 500mM imidazole (elution). Eluted components were further purified by size exclusion chromatography on a Superose 6 Increase 10/300 G/L column (Cytiva 29091596) to select the components that assembled into the correct oligomeric state.

### Array assembly

Refolded SpyTag-fused H-2D(b) bound to the LCMV gp33 peptide (KAVYNFATM) (ST-pMHC) was provided by the NIH Tetramer Facility. Varying molar ratios of ST-pMHC and purified A-SC from 1:3 to 1:192 (ST-pMHC:A-SC) were mixed and incubated overnight at 4C to allow formation of the SpyCatcher-SpyTag bond. The resulting A-pMHC and an unreacted A-SC control were then each mixed with B-c-GFP and diluted to a final concentration of 5uM A-SC and 2.5uM B-c-GFP in TBSG with 500mM imidazole. The final mixture was allowed to assemble overnight at 4C, protected from light. Assembled arrays were separated from free components by centrifugation at 5000xg for 5min. After aspirating the supernatant, the pelleted arrays were resuspended in TBSG with 500mM imidazole to their original volumes before aspirating.

### Cell culture

P14 TCR transgenic Jurkat cells were provided by Matthew Wither and grown in RPMI (Gibco 11835-030) supplemented with 10% FBS (VWR 89510-186), 10mM HEPES (Gibco 15630-080), 1mM sodium pyruvate (Gibco 11360-070), and 1x Pen/Strep/Glu (Gibco 10378-016) (T-cell media). Cells were stored as 500uL aliquots at 5M/mL in T-cell media with 5% DMSO in a liquid nitrogen dewar. Four days prior to imaging, cells were thawed, washed with 2.5mL of T-cell media, plated in 5mL T-cell media in a 6cm dish and incubated for two days at 37C and 5% CO_2_. After two days cells were counted and diluted to 100k cells/mL in T-cell media and incubated for another two days at 37C, 5% CO_2_.

### Jurkat stimulation and staining

All stimulation and staining was performed in a 96-well black-sided glass-bottom plate (MatTek PBK96G-1.5-5-F). The plate was first coated with a mixture of anti-GFP (100x dilution; MBL Life Science 598), anti-LFA1 (10ng/uL; BioLegend 301202), and retronectin (20ng/uL; Takara T100B) in PBS overnight at 4C. After coating, the plate was blocked in 1% casein (Sigma Aldrich C8654) for 1hr at 37C and washed three times with PBS. Each well of the plate was then coated with 50uL of a 2000x dilution of assembled arrays in TBSG with 500mM imidazole. The plate was centrifuged at 100xg for 5min to improve array coating and washed three times with PBS with 0.1% CHAPS, and three times with PBS to remove any arrays and unassembled components not firmly attached. P14 Jurkats were seeded into the plate at 35k cells/well and centrifuged at 100xg for 1min to induce contact with the arrays. After 5 or 10min of incubation at room temperature (including the 1min centrifugation) cells were fixed with BD fixation buffer (BD Biosciences 51-2090KZ) for 30min at room temperature. Fixed cells were permeabilized and blocked with a 1x BD wash/perm buffer (BD Biosciences 51-2091KZ; diluted from 10x in Fc blocking buffer) for 15min at room temperature, protected from light. Cells were then washed twice with 1x BD wash/perm buffer diluted in PBS, and stained overnight at 4C, protected from light, with 50uL/well of PE-anti-pLAT (BD Biosciences 558487; 5x dilution) and AF647-anti-pCD3z (Abcam ab237452; 100x dilution) in 1x BD wash/perm buffer diluted with PBS. Staining solution was removed with two washes with BD perm/wash diluted in PBS and one with PBS. Cells were then stained with 100uL of 5μM CellTrace Violet (Invitrogen C34557) in PBS for 20min at room temperature, protected from light. Excess CellTrace was removed with three washes in PBS, and cells were left in PBS for imaging. All washes and buffer changes were performed with 100uL per well unless otherwise listed and all liquids were added gently to avoid dislodging the cells from the glass surface.

### Imaging

Images of fixed and stained cells were acquired on a Leica DMi8 with a spinning disc confocal, hardware autofocus, and a 63x (0.75 NA) glycerol objective. At each position, a z-stack of 6 slices spaced out by 200nm, centered on the glass surface was acquired. Images of cells (DIC, 100ms exposure), CellTrace (405 Ex, 440/40 Em, 500ms exposure), arrays/GFP (470 Ex; 510/50 Em, 900ms exposure), PE-anti-pLAT (555 Ex, 600/50 Em, 200ms exposure), and AF647-anti-pCD3*ς* (640 Ex, 700/75 Em, 900ms exposure) were acquired at each slice using an LDI-7 from 89North at 50% power.

### Image Processing

Images were processed using a custom MATLAB script to extract information about the CD3ς and LAT phosphorylation in areas where cells came in contact with pMHC arrays. First, a segmentation of the cells was determined from the CellTrace stain, including dimmer regions containing thin regions of cytoplasm in contact with the glass surface. Next, a segmentation of the arrays as determined from the GFP signal in a manner that allows detection of both very bright and very dim arrays. Finally, the total or median fluorescence intensity in each channel (with the local background in a small ring around the array subtracted from each pixel) was calculated for each region where the two segmentations overlapped. As an image-internal control, the shape of each overlapping region was extracted and placed at a random location within a cell in the same image and the same total fluorescence quantification was made in that region.

### GFP Counting

To estimate the number of GFP molecules present in an array from its total intensity, we used particles with known GFP copy number to calibrate our imaging protocol. Nanocages with either 60 or 120 GFP copies attached^46^ were coated onto the glass imaging surface and fixed using the same protocol as the arrays. GFP nanocage images were segmented and quantified using the same custom scripts as the array images and the average total intensity for all segmented nanocages was calculated for each GFP copy number. A linear fit to these data allows interconversion between total GFP fluorescence intensity and GFP copy number (fig. S2).

## Data, Materials, and Software Availability

Code used to run simulations and analyze simulation and imaging data will be uploaded to the Kueh lab github (https://github.com/kuehlab).

## Acknowledgements

We thank members of the Kueh lab, in particular Morgan Bean, for discussion and feedback, Matthew Wither for providing the P14 Jurkat cell line, Tony Cooke (previously with Leica Microsystems) for microscope setup and support, and the NIH Tetramer Core Facility for providing SpyTag-pMHC monomers. This study was funded by an NIH/NIBIB Trailblazer Award (R21EB027327, H.Y.K.), a Bill and Melinda Gates Foundation grant (INV-010680, D.B.), and an NIH NIAID grant (P01AI091580, J.T.G.).

## Author contributions

W.L.W., J.T.G. and H.Y.K. designed the research; W.L.W. performed the simulations; W.L.W. and H.K.Y. performed the experiments; A.J.B. contributed reagents and protocols for array assembly; W.L.W. and H.K.Y. analyzed the data; D.B. provided computational resources; J.T.G. edited the paper; W.L.W. and H.Y.K. wrote the paper.

