## Supplemental Figures for "Proofreading and single-molecule sensitivity in T-cell receptor signaling by condensate nucleation"

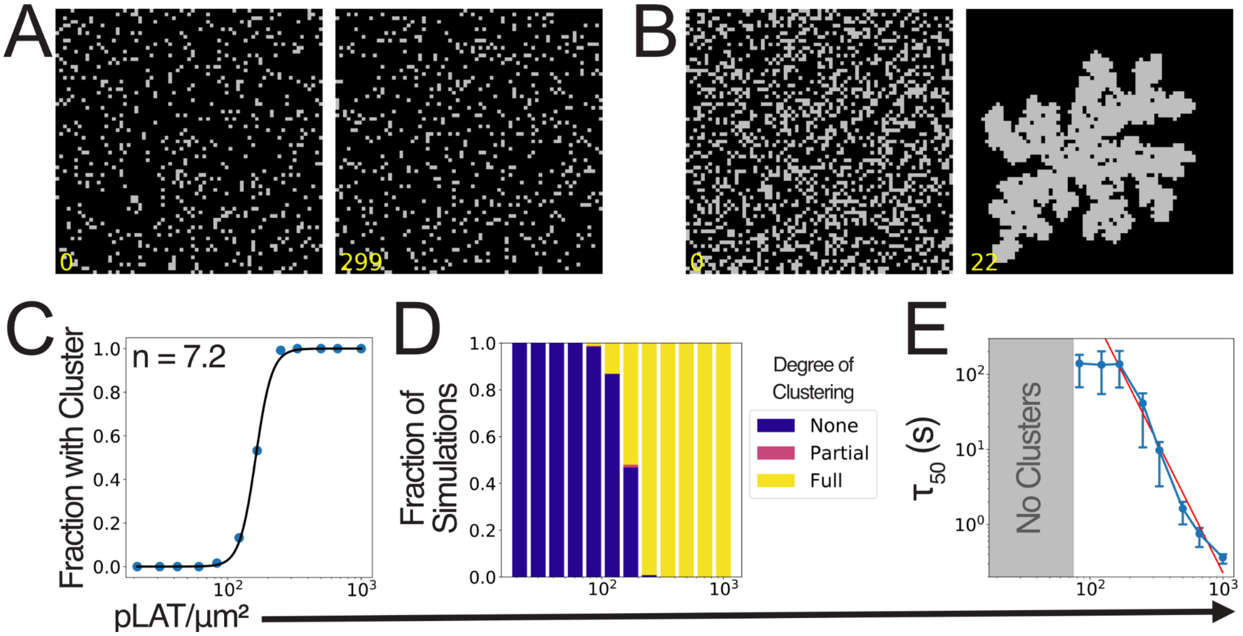


**Figure S1. pLAT molecules condense rapidly above a critical concentration.** Data from 250 replicate simulations without TCR or CD45 activity at varying $p_3$LAT 2D densities. No $p_4$LAT was included in these simulations. A,B) Representative snapshots of simulations with 122pLAT/µm$^2$ (A) or 333pLAT/µm$^2$ (B). Images are 2.13µm square, timestamps are min:sec. C) The fraction of simulations at each pLAT density that achieve a maximum cluster size of at least 50 molecules. Black curve represents the fit to the Hill equation. D) Stacked bar plot showing the proportion simulations that have either full, partial, or no clustering, with >95%, 5%-95%, or <5% of LAT in clusters, respectively. E) The mean and standard deviation of the time until the first cluster for each pLAT density where any clusters form. Red line shows power law fit to the points with pLAT densities achieving at least 50% clustering.

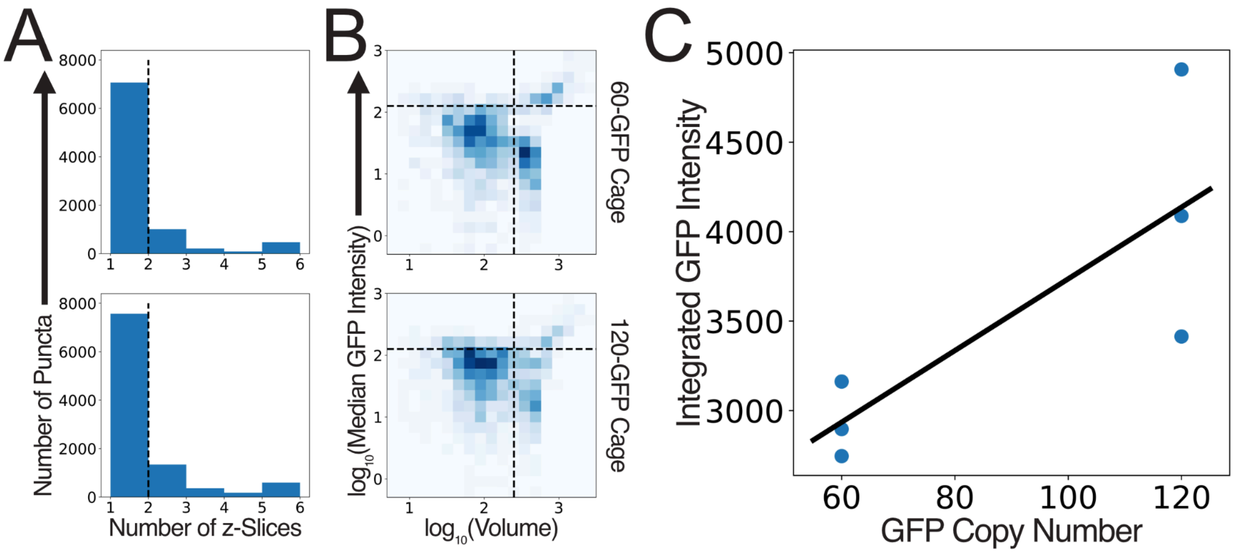


**Figure S2. GFP counting calibration.** A,B) Filtering criteria for inclusion of puncta in GFP counting analysis. A) Histograms showing the distribution of the number of z-slices each nanoparticle spans for the 60-GFP nanoparticle (top) and 120-GFP nanoparticle (bottom). Nanoparticles must be present in at least 2 consecutive z-slices to be included in the analysis. B) 2D histograms showing the joint distribution of the volume (voxels) and median GFP intensity (a.u.) for the 60-GFP nanoparticle (top) and 120-GFP nanoparticle (bottom). Nanoparticles must not exceed 252 voxels or a median intensity of 126 in order to be included in the analysis. C) Linear fit of the median of the integrated GFP intensity distribution of nanoparticles passing the filtering criteria to three replicates at each GFP number.

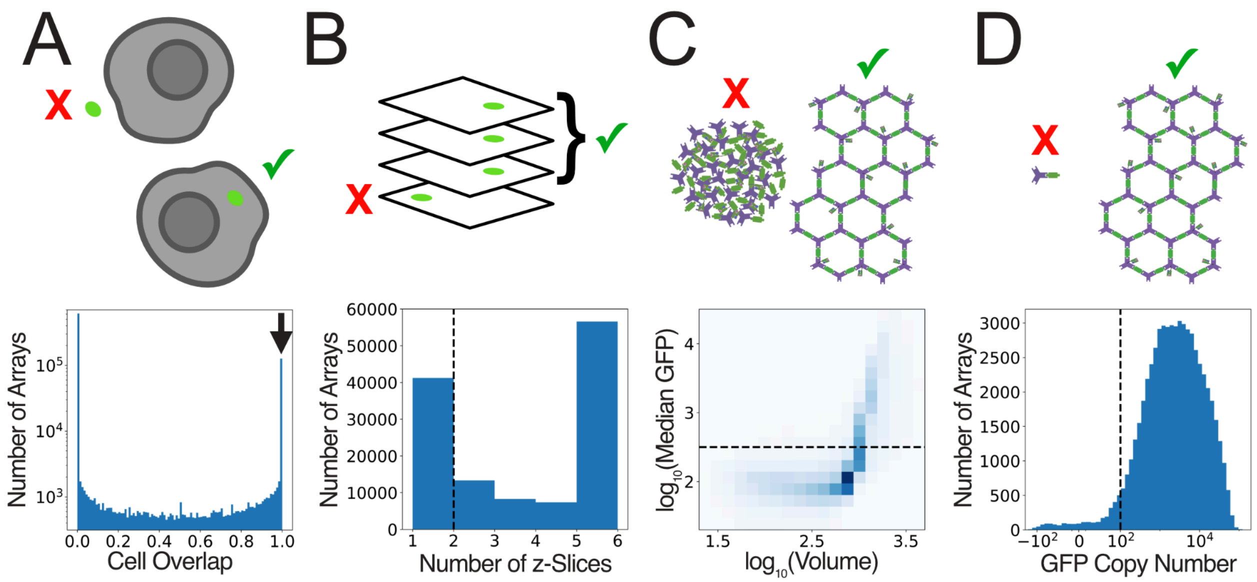


**Figure S3. Filtering removes low quality arrays.** Cartoons and distributions describing the filtering criteria for inclusion of arrays in later analysis. A) Cell contact filter. Histogram shows the distribution of the fraction of the array volume that overlaps with a cell for all arrays. Only arrays with at least 99% overlap are included. B) Z-continuity filter. Histogram shows the distribution of the number of consecutive z-slices the array is present in for all arrays passing the cell contact filter. Arrays must span at least 2 consecutive z-slices to be included. C) Aggregation filter. 2D histogram shows the joint distribution of the volume (voxels) and median GFP intensity (a.u.) of all arrays passing the cell contact and z-continuity filters . Arrays with median GFP greater than 317 are considered to be aggregated and are excluded. D) Size filter. Histogram shows the distribution of GFP copy number for all arrays passing the cell overlap, z-continuity, and aggregation filters. Arrays must have at least 100 GFP copies to be included.

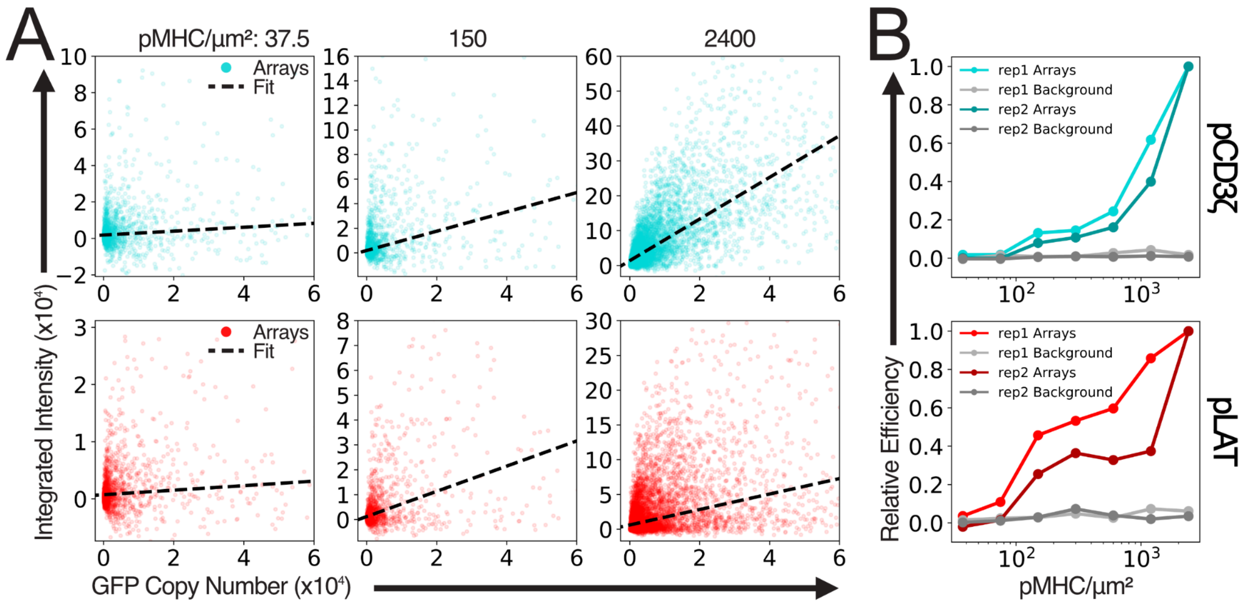


**Figure S4. Total phosphorylation increases with array size and density.** A) Scatterplots showing the background-subtracted integrated pCD3ζ (top) or pLAT (bottom) intensities produced by arrays vs the number of GFP molecules in the array for arrays from three representative pMHC densities. Black dashed lines indicate the line of best fit to the data. B) Line plots showing the relationship between the slope of the fitted lines in (A) and the pMHC density for two biological replicates. Each replicate is independently normalized to the value for the highest pMHC density.

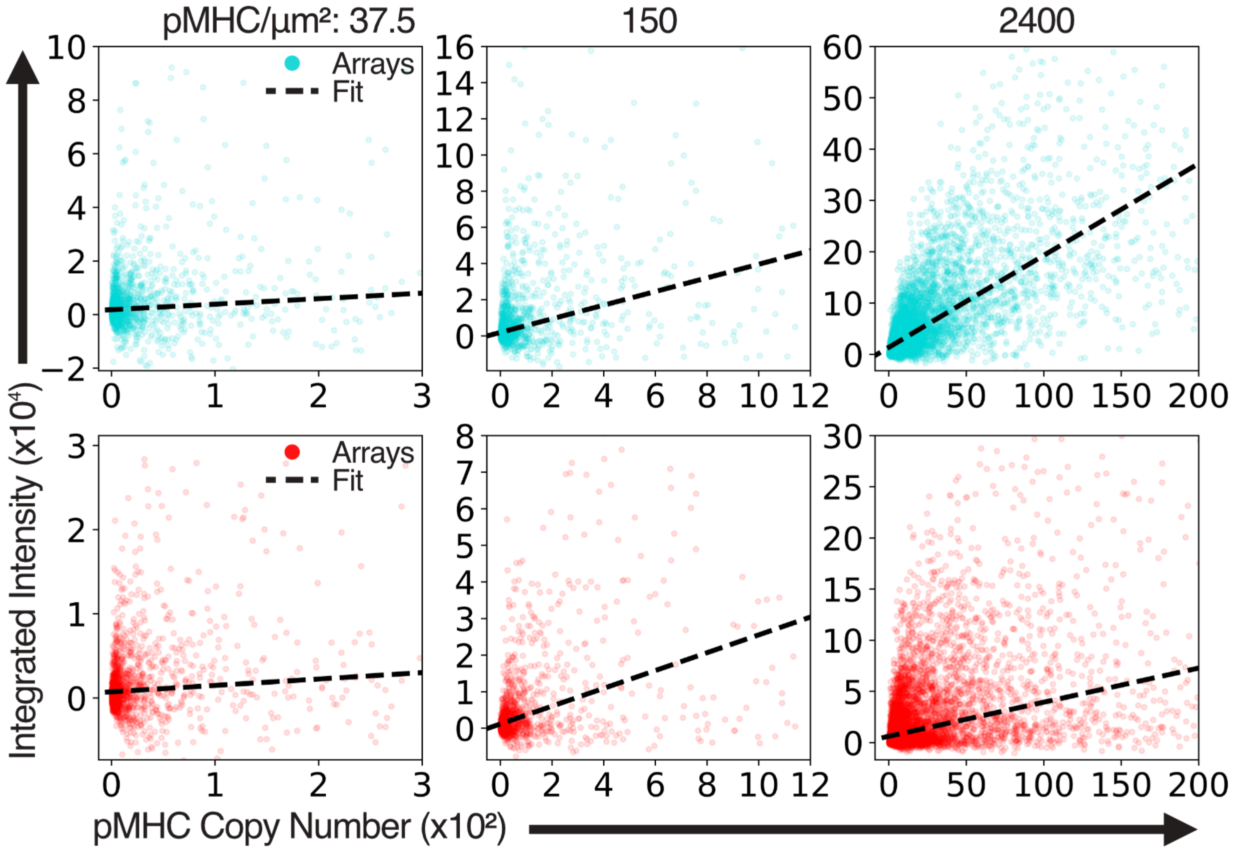


**Figure S5. Phosphorylation at constant array size increases with pMHC density.** Scatterplots showing the background-subtracted integrated pCD3ζ (top) or pLAT (bottom) intensities produced by arrays vs the number of pMHC molecules in the array for arrays from three representative pMHC densities. Black dashed lines indicate the line of best fit to the data.
