## Supplementary figures and images for "Proofreading and single-molecule sensitivity in T-cell receptor signaling by condensate nucleation"

### Movie S1

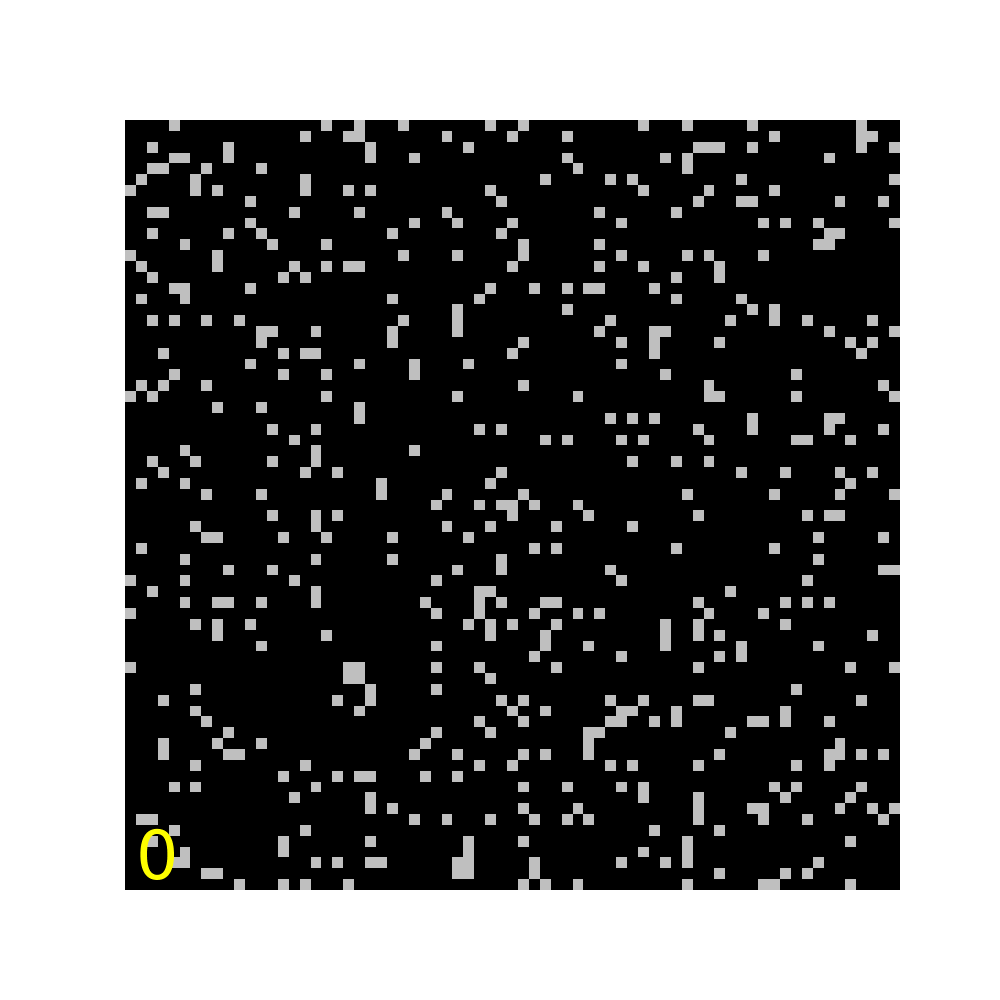

### Movie S2

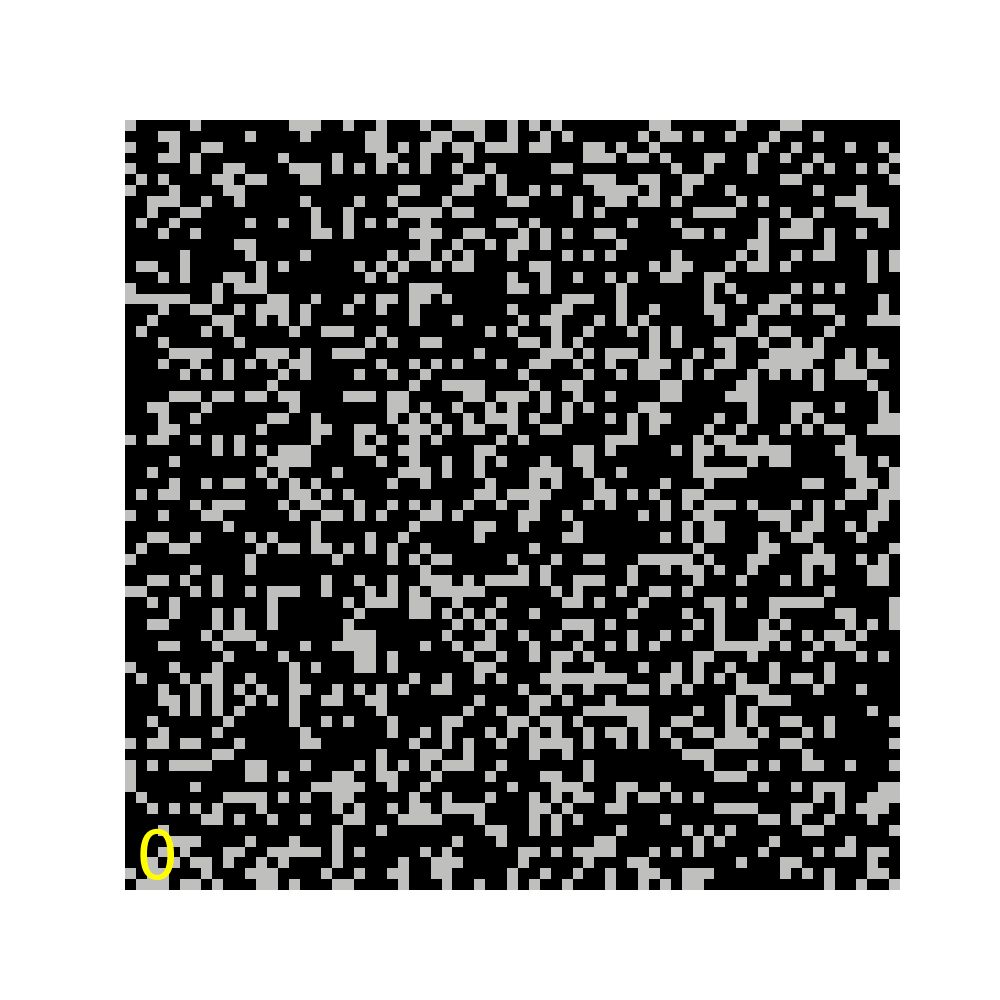

### Movie S3

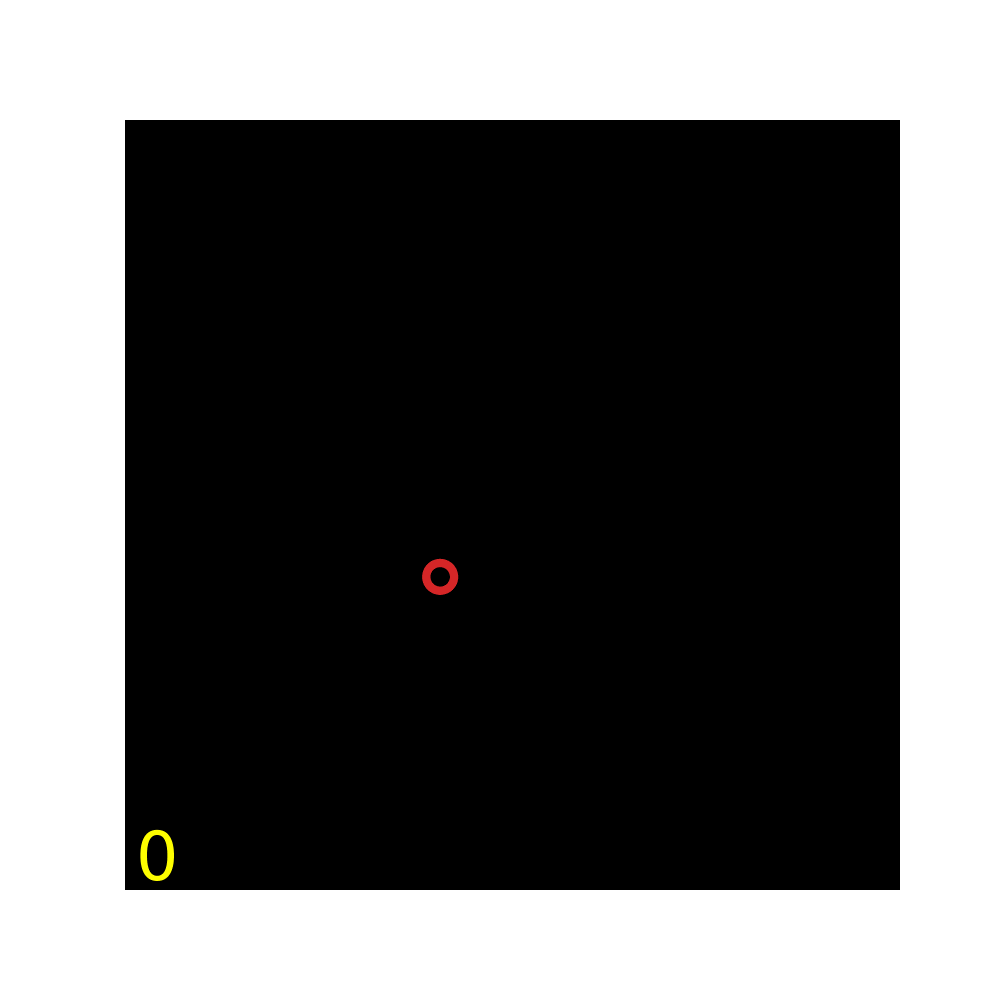

### Movie S4

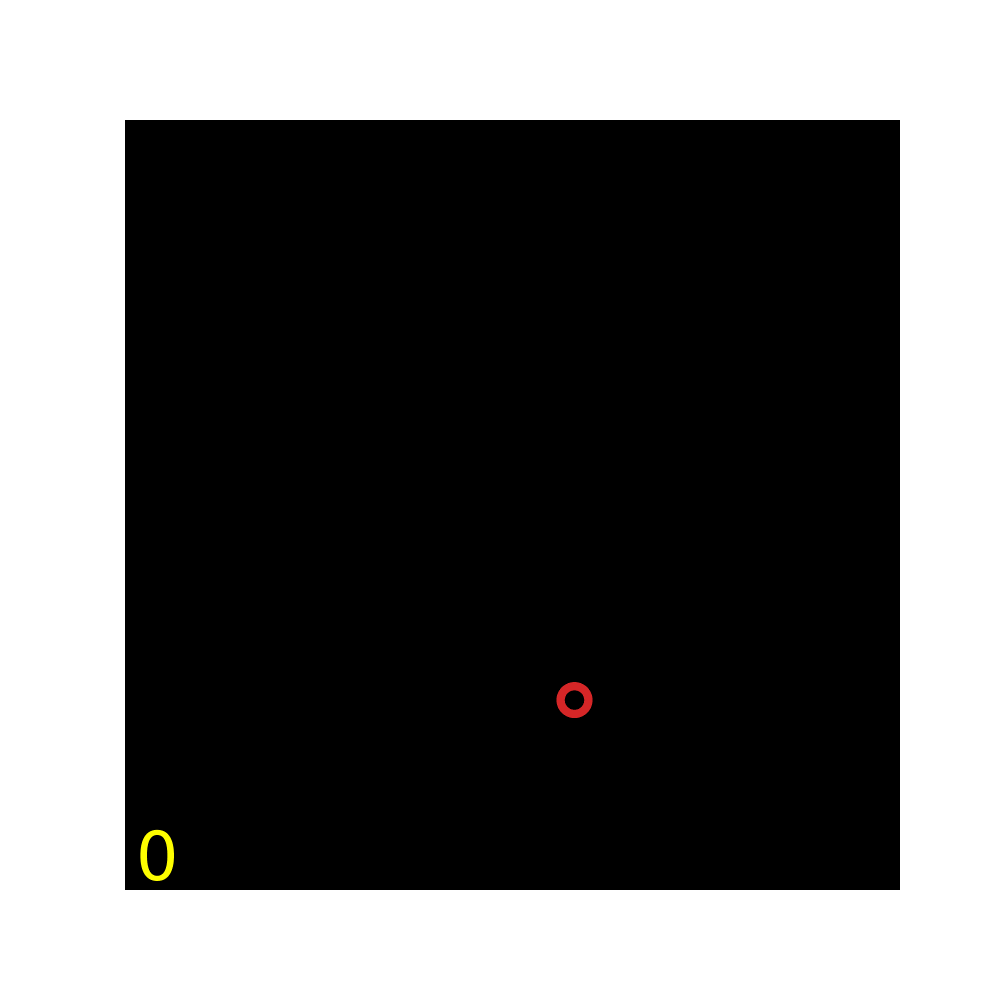

### Movie S5

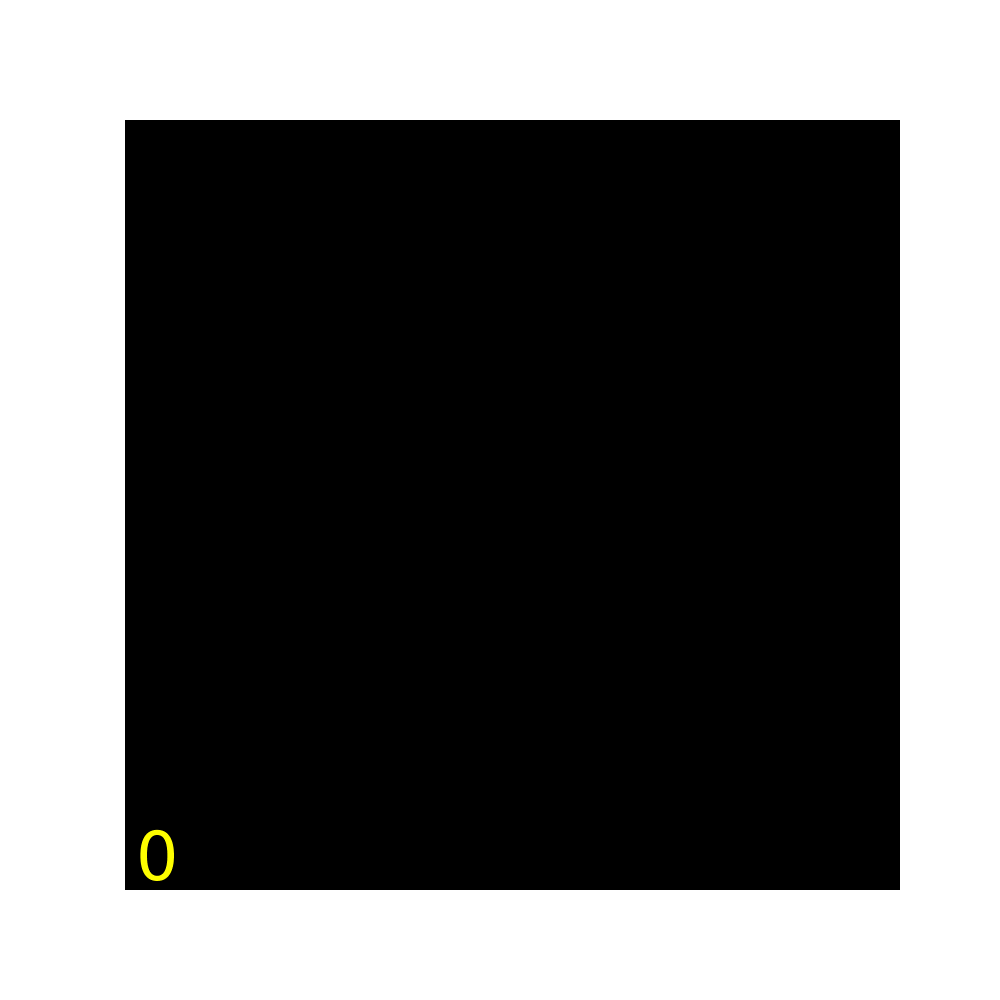

### Movie S6

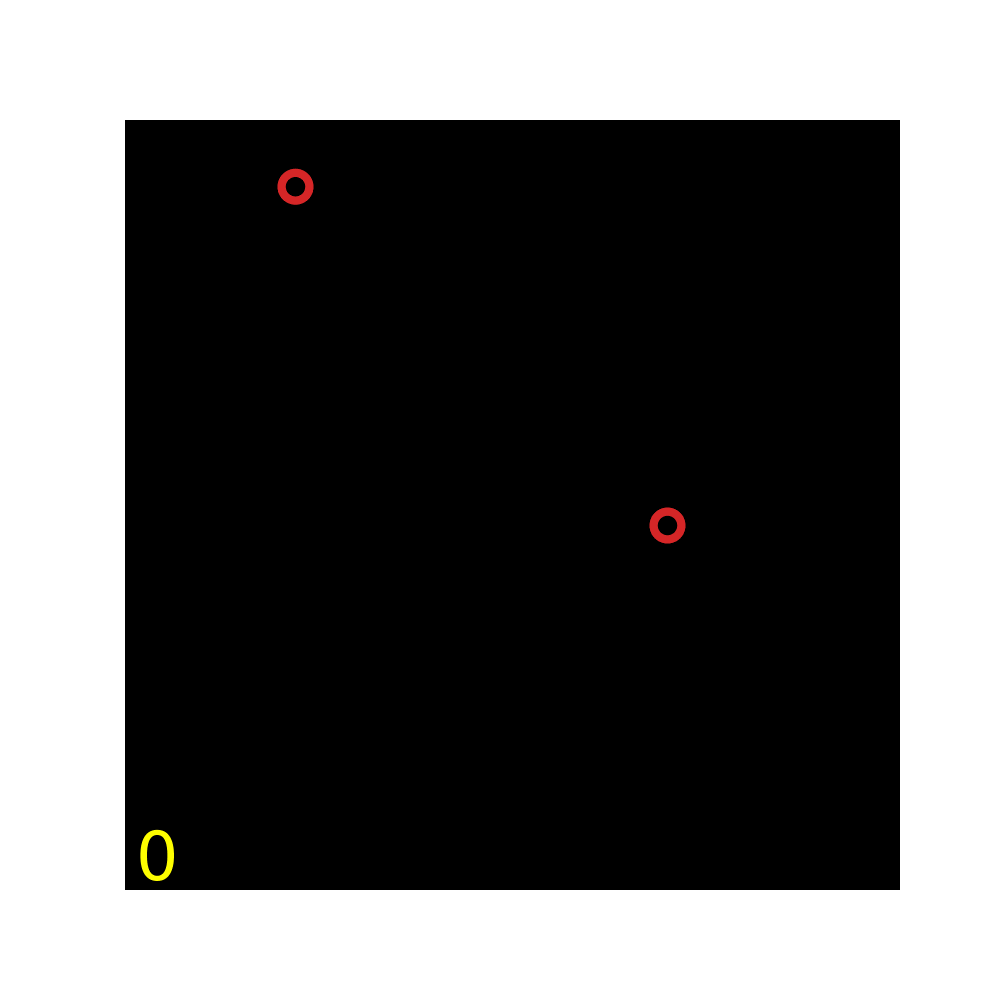

### Movie S7

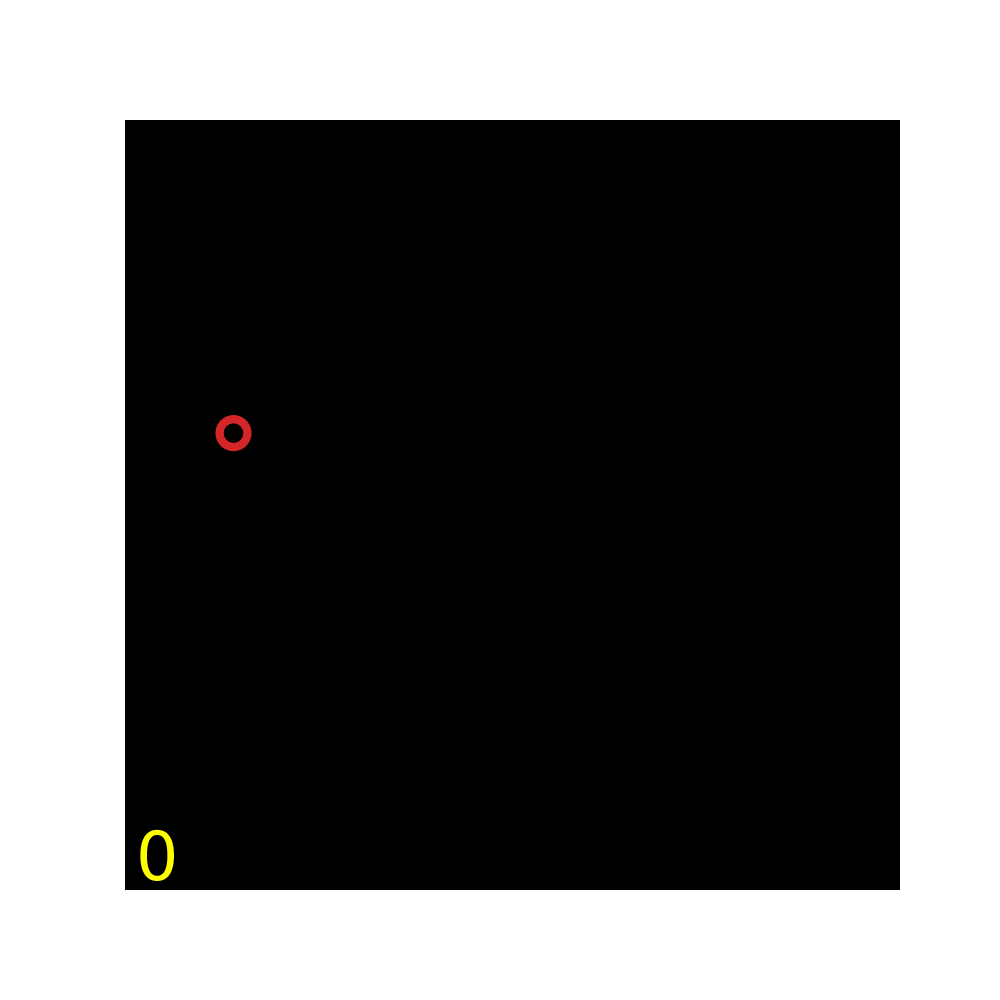

### Movie S8

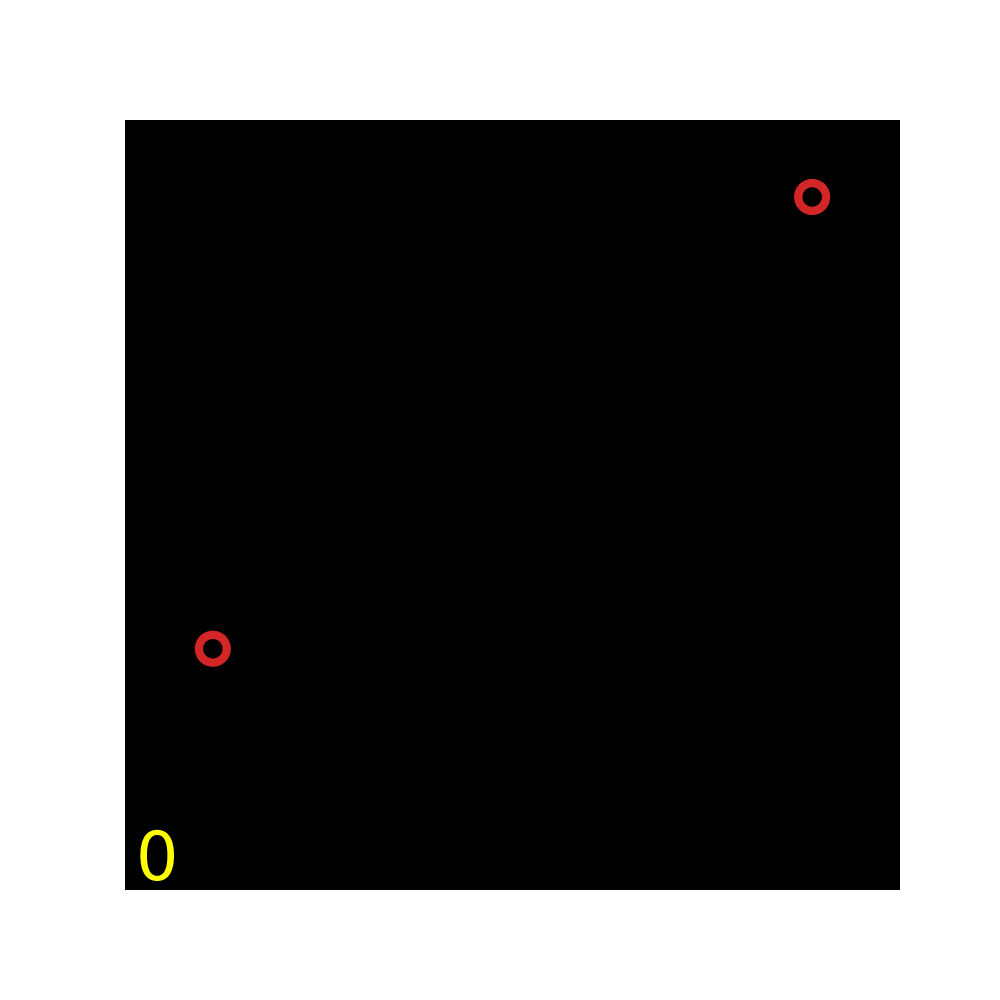

### Movie S9

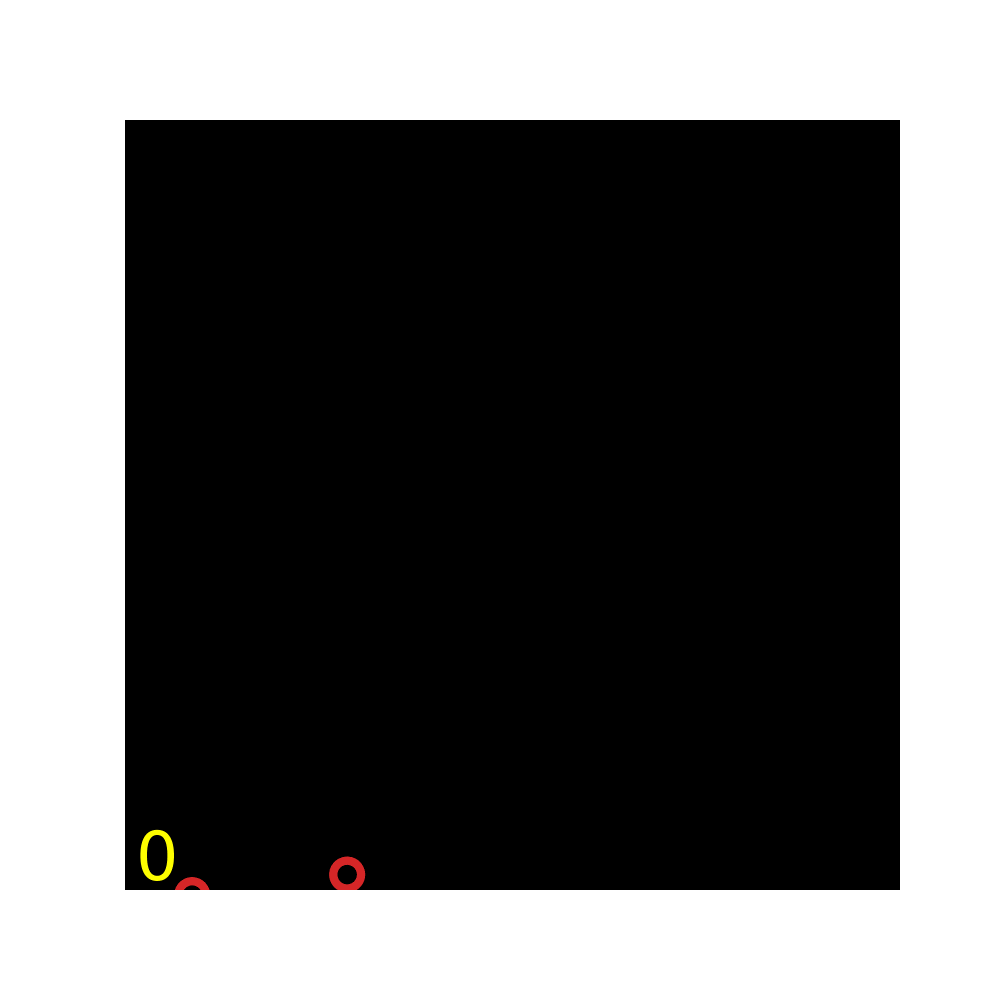

### Movie S10

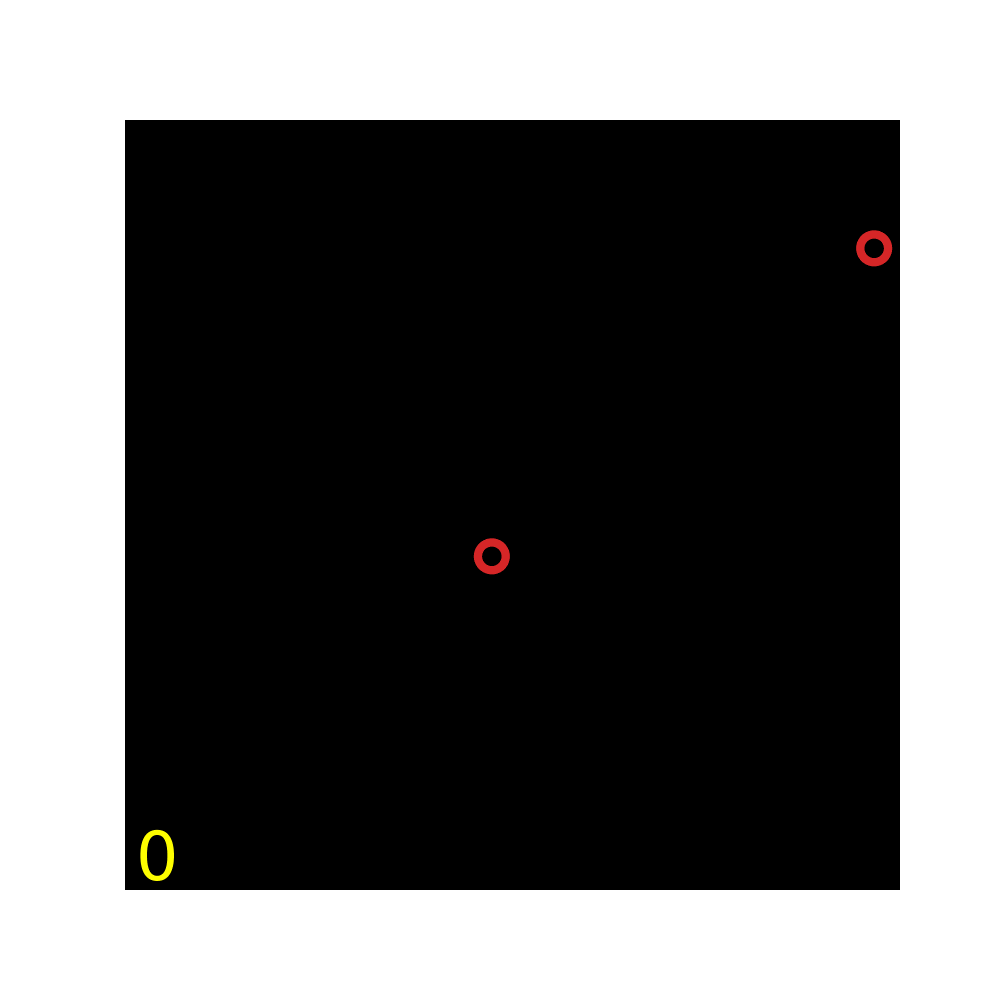

### Movie S11

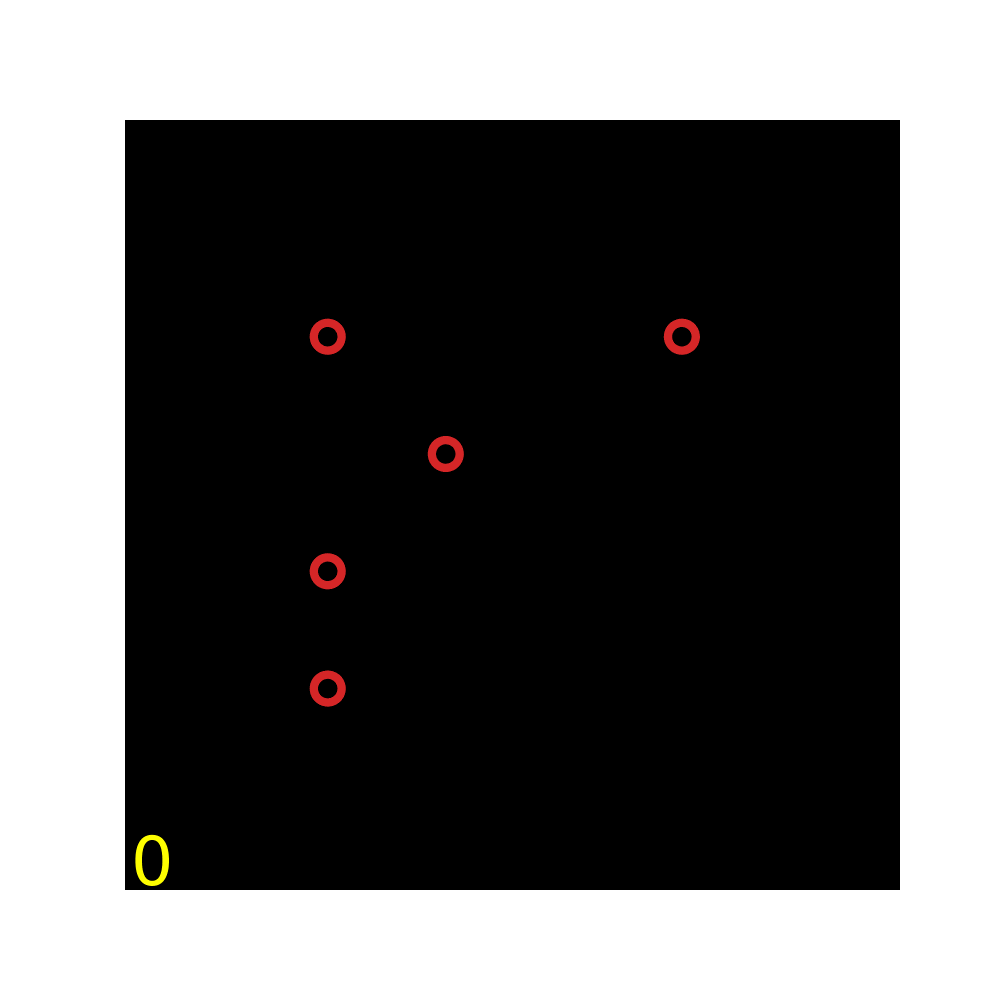

### Movie S12

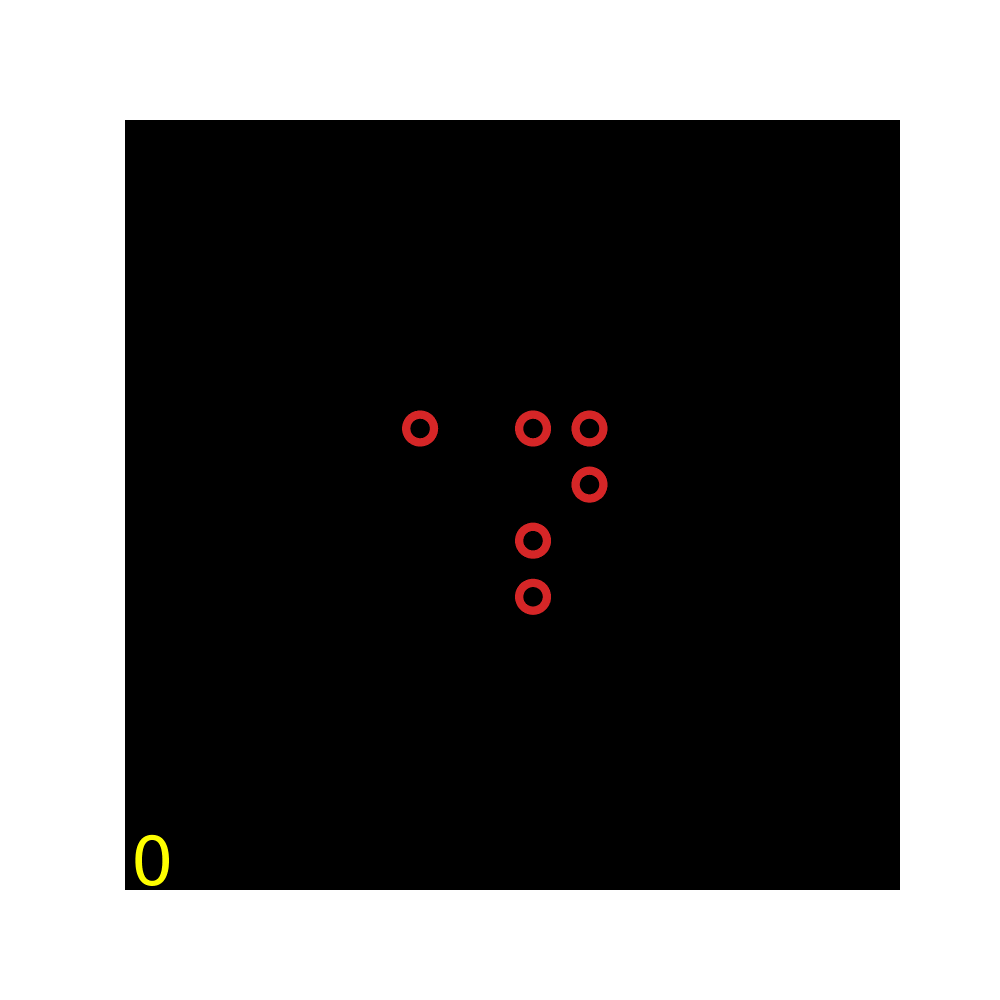

### Movie S13

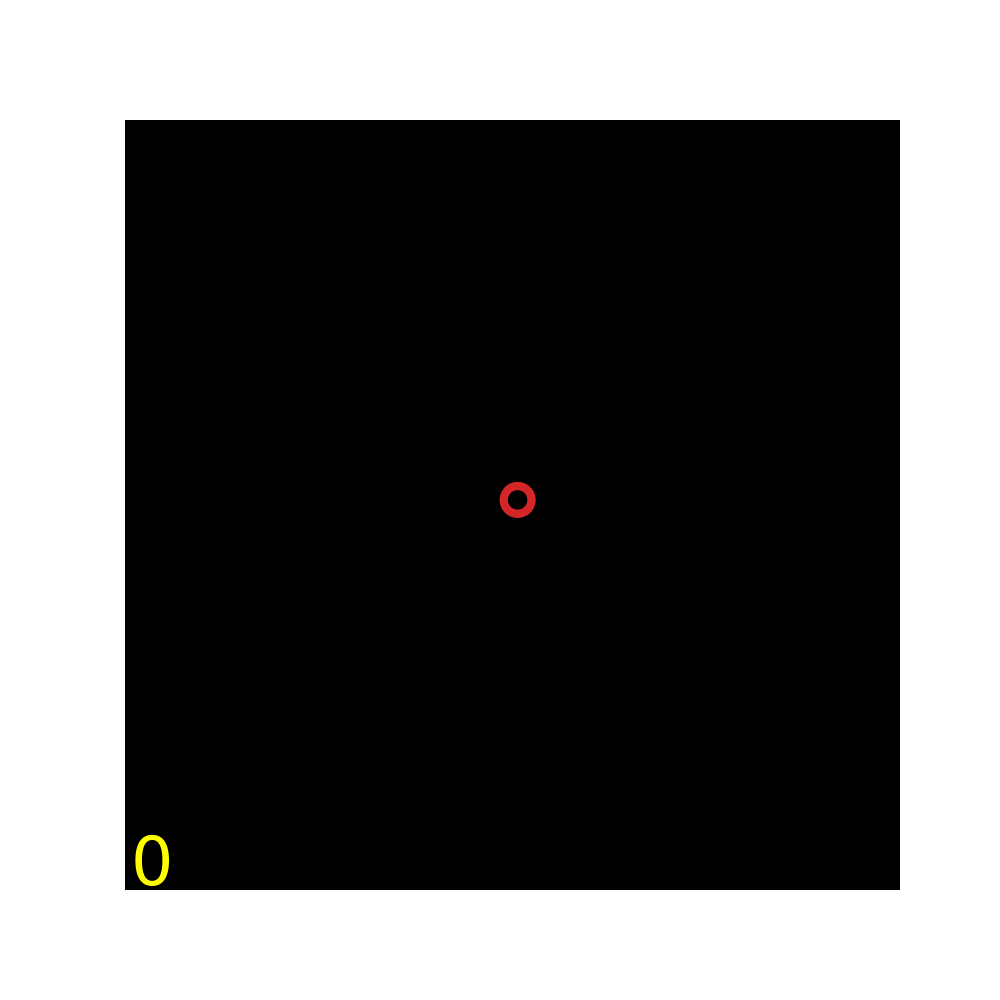

### Movie S14

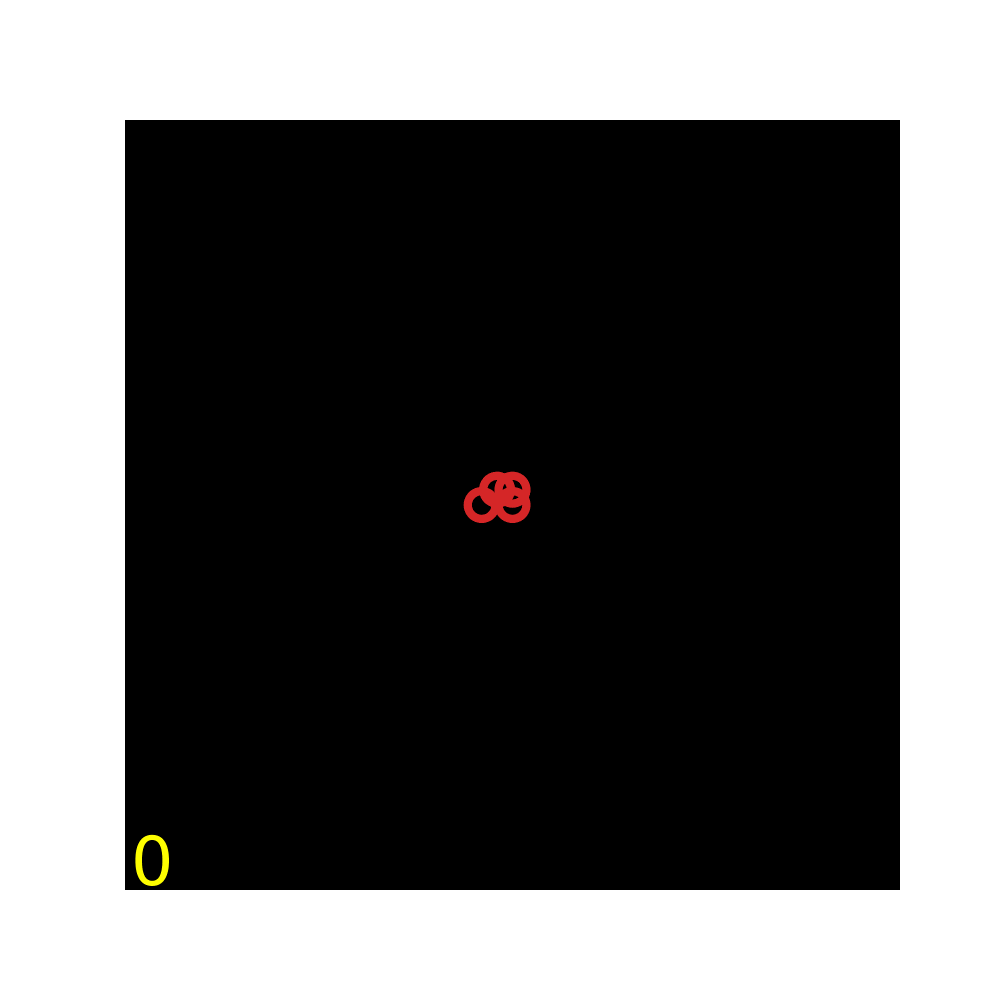

### Movie S15

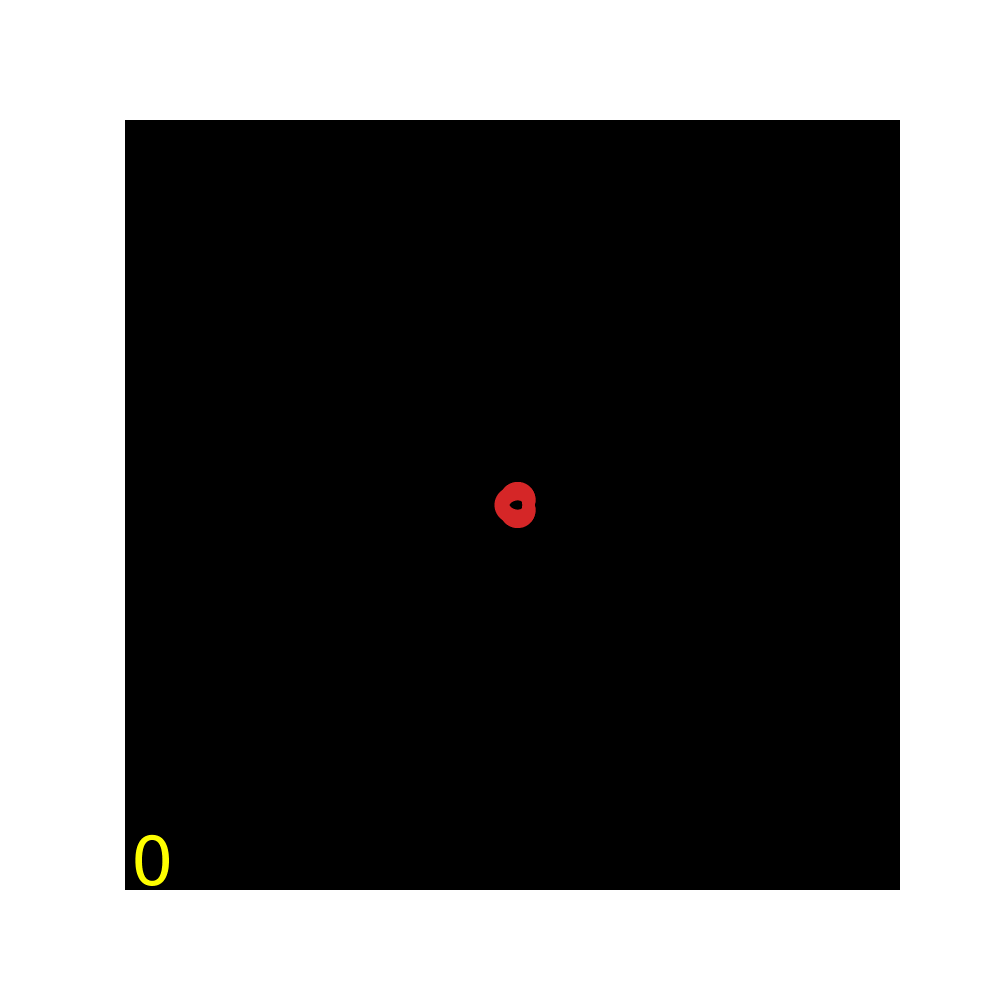
